# Neurod2/6 transcription factors control basal progenitor differentiation and sequential production of neocortical cell subtypes

**DOI:** 10.1101/2025.02.12.637921

**Authors:** Kuo Yan, Andrew G Newman, Svetlana Tutukova, Maria Gavrish, Ingo Bormuth, Olga Bormuth, Victor Tarabykin

## Abstract

Neurod2 and Neurod6 (Neurod2/6) are key transcription factors that promote neuronal differentiation, however, their role in the developing neocortex is not fully understood. Both genes have similar expression patterns during development as well as binding motifs. In order to investigate their role in cortical neurogenesis, we generated a mouse model deficient for both Neurod2/6. Here we demonstrate that differentiation of Tbr2-positive (Tbr2+) basal progenitors (BPs), but not Pax6+ apical progenitors, was severely defected, resulting in ectopically expanded BPs in perinatal Neurod2/6 deficient brains. The sequential fate specification of cortical neurons was also impaired in the absence of Neurod2/6. Ectopic Tbr2+ BPs expressed multiple proliferation markers and were able to self-renew. Olig2+ glial precursors were consequently over-produced in Neurod2/6 deficient brains. Restoration of Neurod2/6 in the double deficient brains downregulated Tbr2 expression, and exhibited substantial rescue effects on defected laminar subtype specification and excessive gliogenesis. Our work indicates that Neurod2/6 regulate BP differentiation and sequential production of cortical cell subtypes via inhibiting Tbr2-dependent genetic program.

## Introduction

During development of the cerebral cortex, all projection neurons originate from neural progenitor cells proliferating in the ventricular and subventricular zones (VZ and SVZ) of the dorsal telencephalon (Subramanian et al., 2017). In the beginning of corticogenesis, this proliferative compartment is a single-layered VZ, with all progenitors contacting the lateral ventricle lumen (Meyer et al., 2000; Homem et al., 2015; Montiel et al., 2016). With development, VZ progenitors elongate their basal process and thereby transform into radial glia cells (RGCs), which play a dual role in development (Malatesta et al., 2003). Like neuroepithelial cells, RGCs function as neuronal progenitors, which are also called apical progenitors (APs) as their cell bodies are lining at the apical surface of neocortex. Additionally, they guide neuronal migration by providing a scaffold for radially migrating neurons through their long basal processes (Rakic, 1972; Levitt and Rakic, 1980).

At the onset of neurogenesis (around E10.5 in mice), some APs undergo asymmetric division and translocate to the basal border of VZ, forming the SVZ. These cells in SVZ are mitotically active, called basal progenitors (BPs), or intermediate progenitors (Noctor et al., 2004 and 2008; Homem et al., 2015; Montiel et al., 2016; Wilsch-Bräuninger et al., 2016). BPs may either generate neurons in one neurogenic division or renew themselves by proliferative mitoses to enrich the neural progenitor pool in SVZ (Noctor et al., 2004; Sessa et al., 2008). SVZ is a critical epicenter for newborn neurons, where they begin to differentiate, extend axons, and start radial migration along radial glia fibers (Bayer and Altman, 1990). SVZ is one of the hotspots in the evolution of the neocortex. It is relatively small in nonplacental mammals and rodents, and comparable in size with that of the VZ. SVZ size increases proportionally to the increment of the neocortex thickness in primates and humans, and significantly overtakes the size of VZ. Respectively, the proportion of proliferating cells in the SVZ of primates is higher than in the VZ.

Tbr2 is a transcription factor (TF) expressed in the SVZ to control BP genesis and proliferation ability. Tbr2 over-expression converts APs to BPs, and prevents BP cell cycle withdrawal (Sessa et al., 2008 and 2017), but it is not known what molecular cascades inactivate Tbr2 in order to initiate pyramidal neuron differentiation. Two members of NeuroD family TFs (Neurod2/6) are potential candidates for this role as their expression arises at the basal border of SVZ where newborn neurons exit cell cycles. While Tbr2 expression vanishes in neurons leaving the SVZ, Neurod2/6 expression is immediately onset in migrating and postmigratory neurons (Wu et al., 2005; Bormuth et al., 2013).

In this work, we investigated the role of NeuroD family TFs in neocortical differentiation and cell fate specification. As both genes have very similar expression patterns and similar targets sets (Bormuth et al., 2013), we inactivated both of them to avoid functional redundancy. This resulted in severe disruption of neocortical differentiation and caused excessive production of Tbr2-positive (Tbr2+) BPs. Additionally, cortical cell subtype specification was also disrupted in the absence of Neurod2/6. Our data indicate that NeuroD2/6 control onset of neuronal differentiation by suppressing Tbr2-dependent transcription program.

## Materials and Methods

### Mice

The transgenic mice (Neurod6-Cre and Neurod2-Null) and genotyping methodology used in this study were previously described (Goebbels et al., 2006; Bormuth et al., 2013; Yan et al., 2023). The Neurod2/6 double deficient (DKO) mice we used in this report are Neurod2^NeoR/NeoR^; Neurod6^Cre/Cre^, while the control mice are Neurod2^Wt/NeoR^; Neurod6^Cre/Cre^. Generation of NeuroD2/6 DKO mice/embryos: a male and a female control mouse were bred to produce a litter of mixed genotypes (25% incidence of NeuroD2/6 mice/embryos, while 50% incidence of controls). The day of plug discovery was termed as embryonic day 0.5 (E0.5). Male and female embryos were not discriminated in this study. All animal experiments conform to German regulatory standards, and were approved/licensed by the local responsible authority (Landesamt für Gesundheit und Soziales Berlin, LaGeSo, Berlin).

### *In utero* electroporation (IUE) and constructs

IUE was performed as previously described (Yan et al., 2023). Briefly, the anesthetized pregnant mice were kept laid down on a heating pad with constant inhalation of isoflurane (mixed with oxygen). The subcutaneous administration of pain killers (such as temgesic) was applied before a surgery started. An incision of ∼15mm was made on the fur and skin along the abdomen midline after cleaned by 70% ethanol. The plasmids (1 µg/µl, mixed with fast green dye 1:20, 000) were loaded in a fine glass capillary and enforced by a vacuum pico-pump into either of the cerebral lateral ventricles. Electrodes were positioned biauricular (anode at the side of injection). Electrodes were positioned biauricular (anode at the side of injection). Electroporation was achieved by an electroporator (CUY21, Sonidel) with setup: 6 times pulses, 35 V voltage, 50 ms pulse duration and 1 sec interval time. The mice on surgeries were monitored every day as for their health till being sacrificed. Our lab generated a Cre-dependent conditionally bicistronic expression vector (pCAG-FPF-GFP) (Parthasarathy et al., 2014; Yan et al., 2023), which the open reading frames (ORFs) of targeted genes (such as Neurod2 and Neurod6) with their original Kozak sequences were subcloned into for IUE experiments. Tbr2 ORF was subcloned into a constitutive expression vector (pCAGIG, Addgene #11159) for over-expression by IUE. All cloned genes were verified by sequencing. Cloning primers are listed in Tab. S1.

### Immunofluorescence (IF)

*Day One*: the brain cryo-sections (16 µm in thickness) were post-fixed in 4% paraformaldehyde (PFA, dissolved in 1x PBS) for 15 min followed by twice washes in 1x PBS (10 min each, the same below or otherwise indicated). Subsequently, the sections were incubated in blocking solution (BS, 1x PBS containing 0.5% Triton X-100, 2% BSA and 10% horse serum) at room temperature (RT) for 1 hour, followed by primary antibodies in BS at 4°C over night (O/N). *Day Two*: the sections were first washed 3 times in 1x PBS and then incubated with secondary antibodies in BS for 1.5 hours. After additional 3 times washes in 1x PBS, the slides were mounted in DAKO anti-artifact medium. As for some antibodies that require antigen unmasking (like BrdU antibody), the sections were immerged in unmasking solution (VectorLabs, H-3300) and constantly heated for 10 min by microwave before blocking on *Day One*, followed by cooling to RT. All primary antibodies used in this study are listed in Tab. S2. Fluorescent secondary antibodies are all from Jackson ImmunoResearch (all were raised in donkey, reconstituted according to manual, and diluted 1:500 in use).

### I*n situ* hybridization (ISH)

ISH was performed as previously reported (Bormuth et al., 2013; Yan et al., 2023). Briefly, *Day One*: the brain cryo-sections were dried in vacuum for 30 min and fixed in 4% PFA dissolved in DEPC-treated 1x PBS (DPBS) for 15 min followed by twice washes in DPBS. Subsequently, the sections were incubated in proteinase K solution (20 mM Tris pH7.5, 1 mM EDTA pH8.0, 20 µg/ml proteinase K) for 2.5 min followed by one wash in DPBS containing 0.2% glycine. The sections were then post-fixed in 4% PFA containing 0.2% glutaraldehyde in DPBS for 15 min followed by twice washes in DPBS, pre-hybridized in hybridization buffer (HB, 50% deionized formamide, 5x SSC, 1% blocking reagent (Roche, 11096176001), 5 mM EDTA, 0.1% Tween20, 0.1% CHAPS, 0.1 mg/ml Heparin, 100 µg/ml yeast RNA) at 65°C for 2 hours, and eventually hybridized with the denatured probes (the probes were heated at 90°C for 5 min before use) at 68°C O/N. *Day Two*: the sections were washed once in 2x SSC pH4.5, and then incubated in RNase solution (0.5 M NaCl, 10 mM Tris pH8.0, 20 µg/ml RNase A) for 30 min at 37°C.

Subsequently, the sections were washed stringently 3 times in 50% formamide/2x SSC pH4.5 (30 min each) at 63°C, followed by 3 times washes in KTBT buffer (50 mM Tris pH7.5, 150 mM NaCl, 10 mM KCl, 1% Triton X-100) at RT. Then the tissues were blocked in ISH BS (KTBT containing 20% sheep serum) for 2 hours, followed by incubation with anti-digoxigenin antibody (alkaline phosphatase (AP)-conjugated, Roche, 1:1500) in BS at 4°C O/N. *Day Three*: the sections were washed 4 times in KTBT (30 min each), washed twice in NTMT buffer (100 mM Tris pH9.5, 100 mM NaCl, 50 mM MgCl_2_, 0.1% Tween20), and were eventually incubated in NTMT containing NBT/BCIP (1:50, AP substrates, Roche). The staining was monitored hourly until the signals showed up. The stained sections were subject to an ascending alcohol series (50% - 100%), incubated in clearing solution (benzyl alcohol:benzyl benzoate = 1:2) for 5 min, and finally mounted in Permount mounting media (Thermo Fisher Scientific). All home-made solutions used on *Day One* were DEPC-treated. Primers used for generation of targeted ISH probes are listed in Tab. S1.

### Bromodeoxyuridine (BrdU) pulse chase

300 μl BrdU solution (10 mg/ml BrdU dissolved in 1xPBS) was injected intraperitoneally into a mouse with a sterile needle. The time points and embryonic stages (E12.5 – E17.5) of injections were labelled for each mouse. The injection and sacrifice time points at various stages were tightly controlled (± 0.5 hour). The mouse brains carrying BrdU incorporation were dissected and fixed in 4% PFA either 24 hours later (cell cycle exit analysis) or at P0 (Tbr2+ cell birthdating analysis). The antigen exposure of BrdU during IF was achieved by constant heating as mentioned in IF.

### Neurite branchy analysis

The cortical cells labeled by electroporated membrane-anchored GFP (or bicistronic Tbr2-GFP) were imaged using Leica SL confocal microscope (63x magnification, 2x zoom, 1 μm spaced z-stacks). The neurite complexity of imaged cells was quantified by Sholl analysis (soma as centroid) using the Neuroanatomy plugin of ImageJ (Fiji).

### Statistics

Cell number or ratio quantifications between control and DKO brains were analyzed with two-tailed *Student’s* t-test. The statistics for dynamic ratios of Tbr2/BrdU double positive cells relative to all BrdU+ cells (Fig. 2E) was analyzed with two-way *Anova* test. The values for Sholl analysis of neurite arborization (Fig. S7) was analyzed by normality and lognormality test by Prism (Milosević, 2007). Values of GFP+ neurons follow (n = 30) normal distribution (*p* = 0.024) and values of Tbr2-expressing neurons (n = 25) is close to normal distribution (*p* = 0.074). The statistics of Sholl analysis was analyzed with unpaired t-test. Bar charts present mean values ± standard deviation (SD). Significance: p<0.0001, ****; p<0.001, ***; p<0.01, **; p<0.05, *; p>=0.05, ns.

## Results

### Neurod2/6 deficiency causes expansion of Tbr2+ basal progenitors

Previously we reported the defect of axon growth and agenesis of corpus callosum in Neurod2/6 double deficient (DKO) mice (Bormuth et al., 2013; Yan et al., 2023). We also noticed that simultaneous inactivation of Neurod2 and Neurod6 in the developing neocortex caused anomalies in numbers and positioning of neocortical neurons (Bormuth et al., 2013). In order to investigate potential roles of Neurod2/6 genes in cell fate specification and differentiation, we analyzed the neocortical progenitor cohorts. We first asked if basal progenitors (BPs) are affected when Neurod2/6 are deleted. We thus performed *in situ* hybridization (ISH) and immunofluorescence (IF) for Tbr2, the major BP determinant (Arnold et al., 2008; Sessa et al., 2008), at different stages of corticogenesis. Strikingly, we found progressive accumulation of Tbr2+ BPs in the SVZ and intermediate zone (IZ) of developing DKO brains (Fig. 1A): the number of Tbr2+ BPs in DKO brains is similar with that of controls at E13.5 but significantly increased by 15.92% ± 8.49% at E15.5 (*p* = 0.029, *), and almost triple of the number of BPs in controls perinatally (*p* = 0.0023, **) (Fig. 1B). Consistently, the expression of BP marker genes, such as Scrt2, Svet1 and Brn1 (Paul et al., 2014; Tarabykin et al., 2001; Sugitani et al., 2002), was also expanded in the SVZ/IZ of DKO brains (Fig. S1).

**Figure 1.**
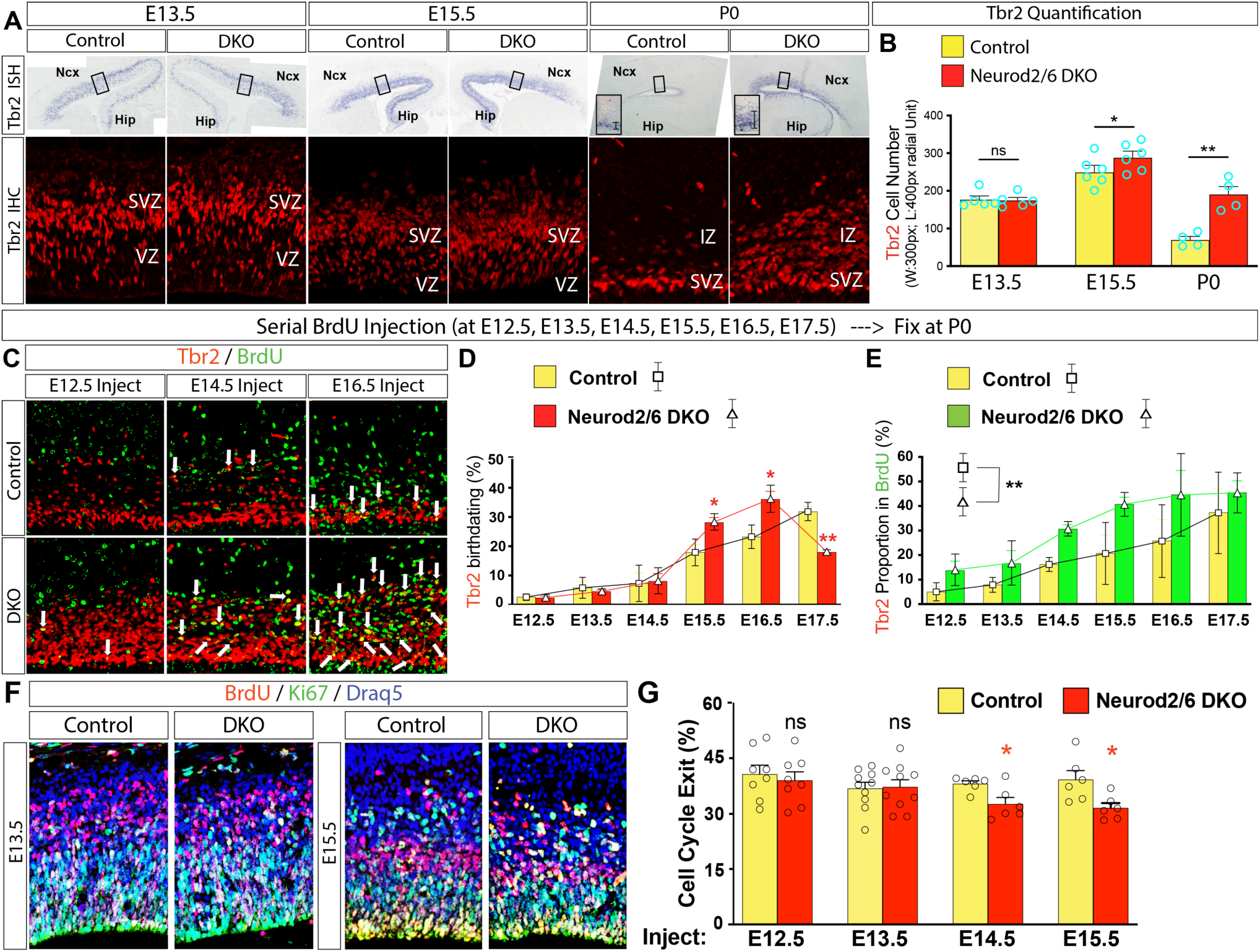
Tbr2+ basal progenitors are ectopically upregulated in Neurod2/6 DKO brains. **(A, B)** *In situ* hybridization (ISH, upper panel) and IF (lower panel) for Tbr2 on control and DKO brain sections at the age of E13.5, E15.5 and P0 shows the progressive emergence of ectopic Tbr2+ basal progenitors (BPs) in the IZ of DKO brains (**A**). (**B**) Cell quantification shows the number of Tbr2+ cells in DKO brains is similar with that of control at E13.5 (*p* = 0.87, ns; n = 5), and is mildly elevated by 15.92% ± 8.49% at E15.5 (*p* = 0.029, *; n = 6). Notably, the number of Tbr2+ cells in DKO brains is increased by 1.73 ± 0.33 folds than that of control at P0 (*p* = 0.0023, **; n = 4). **(C-E)** Mouse brains carrying BrdU injected at serial stages of cortical neurogenesis (E12.5 to E17.5) were fixed at P0 and analyzed by IF for BrdU and Tbr2. Each arrowhead indicates one or one cluster of BrdU/Tbr2 double positive cells (**C**). (**D**) Birthdating analysis for Tbr2+ cells by proportional quantification in control and DKO brains at P0. The production of Tbr2+ cells at P0 displays progressively proportional increment along developmental timing in control brains, however, the proportion of Tbr2+ cell production is substantially elevated at E15.5 (*p* = 0.029, *; n = 3) and E16.5 (*p* = 0.022, *; n = 3), but steeply reduced at E17.5 (*p* = 0.0017, **; n = 3) in DKO brains. Two-way *Anova* analysis is not applied in this scenario because of the existing phenotypes and different denominators. (**E**) Proportional quantification of BrdU/Tbr2 double positive cells relative to total BrdU+ cells in all cortical layers analyzed by two-way *Anova* statistics. There is a significantly larger proportion of BrdU+/Tbr2+ cells accumulated in the SVZ/IZ of DKO brains (F(DFn, DFd) = F(5, 10) = 5.59, *p* = 0.0086, **; each stage n = 3). **(F, G)** Cell cycle exit analysis at serial developmental stages of corticogenesis (BrdU injected at E12.5 to E15.5). The control and DKO brains were fixed 24 hours after BrdU injection and analyzed by IF for BrdU and Ki67 (Draq5 nuclei staining as counter staining, **F**). The proportions of progenitors quitting cell cycles after one day are counted as the numbers of BrdU+/Ki67-cells relative to total BrdU+ cell numbers (**G**). The progenitor proportions exiting cell cycles in DKO cortices are similar with those of controls at E12.5 (*p* = 0.62, ns; n = 8) and E13.5 (*p* = 0.87, ns; n = 10), however, the proportions are significantly reduced in DKO brains in comparison to those of control brains at E14.5 (by 14.48% ± 5.10%, *p* = 0.018, *; n = 6) and E15.5 (by 19.55% ± 7.11%, *p* = 0.021, *; n = 6).

In order to further study the origin of excessive Tbr2+ BPs in DKO brains, we carried out BrdU chase analysis. Pregnant mice were injected with thymidine analog BrdU at serial stages of cortical development (every 24 hours from E12.5 to E17.5). BrdU/Tbr2 colocalization in the neocortex was tested at P0 (Fig. 1C). In such experiments, BrdU will be incorporated into the cells in S phase at the time of injection. On the other hand, cells that contain high levels of BrdU at P0, are the cells that exited mitotic cycle shortly after BrdU injection. The ratios of Tbr2/BrdU double positive (Tbr2+/BrdU+) cells relative to all Tbr2+ BPs indicate that the majority of ectopic Tbr2+ BPs in DKO brains were born after E14.5 (Fig. 1D). Moreover, we quantified the ratios of Tbr2+/BrdU+ cells born from E12.5 till E17.5 relative to all BrdU+ cells at P0. This number reflects the fraction of cells that left mitotic cycle at a defined stage of corticogenesis that maintained Tbr2 expression at P0. In control brains, a very small fraction (less than 5%) of cells that were born at E12.5 still retained Tbr2 and were located in the SVZ/IZ. In DKO brains, such a fraction was almost doubled. This trend was also observed for the newly born cells at every tested stage of development (Fig. 1E). These data indicate that inactivation of Neurod2/6 in the developing neocortex causes arrest of neuronal differentiation, retention of postmitotic cells in the SVZ/IZ and inability to inactivate Tbr2-dependent genetic program.

Next we asked whether Neurod2/6 deficiency could influence cell cycle exit and onset of neuronal differentiation. Here we utilized 24 h BrdU chase. In these experiments, BrdU was injected at four stages: E12.5 till E15.5. The proportions of progenitors quitting cell cycles after one day were counted as the numbers of BrdU+/Ki67-cells relative to total BrdU+ cell numbers (Fig. 1F). The progenitor proportions exiting cell cycle in DKO cortices are similar with those of controls at E12.5 (*p* = 0.62, ns) and E13.5 (*p* = 0.87, ns), however, the proportions were significantly decreased in DKO brains in comparison to those of control brains at E14.5 (by 14.48% ± 5.10%; *p* = 0.018, *) and E15.5 (by 19.55% ± 7.11%; *p* = 0.021, *) (Fig. 1G). As Neurod2/6 are expressed in Tbr2+ BPs but not in Pax6+ apical progenitors (see below), these data indicate that it is a fraction of Tbr2+ cells that retain cycling in DKO brains.

### Neocortical cell type specification is disrupted in Neurod2/6 deficient mice

We reasoned if Neurod2/6 deficiency causes such a dramatic change in the cortical differentiation program, so that many cortical cells retain Tbr2 expression and do not commence radial migration, it might cause alterations in neocortical cytoarchitecture. We quantified the proportions of different cell types in the developing neocortex. We found that the proportion of Satb2+ upper layer (UL) neurons was increased by 25.11% ± 7.18% (*p* = 0.0081, **) while the proportion of Ctip2+ deeper layer (DL) neurons was decreased by 38.20% ± 6.89% (*p* = 0.00050, ***) in DKO brains compared with control brains at E15.5. Satb2 is an UL neuron specifying TF that acts by suppressing Ctip2 expression (Britanova et al., 2006 and 2008). However, there are a subset of transient Satb2+/Ctip2+ neurons in the developing neocortex. The number of such Satb2+/Ctip2+ cells was almost doubled in DKO brains (*p* = 0.026, *) (Fig. 2A, B). IF for other layer specific markers such as Brn2 (UL, Dominguez., et al., 2013; Sugitani et al., 2002) and Tbr1 (DL, Hevner et al., 2001) at E15.5 confirmed the shift in cell fate specification (Fig. 2C). It turned out that the proportion of Brn2+ UL neurons was significantly increased by 27.17% ± 10.35% (*p* = 0.030, *) while the proportion of Tbr1+ DL neurons was decreased by 29.50% ± 7.39% (*p* = 0.0040, **) in the cortical plate (CP) of DKO brains (Fig. 2D, 2E). Additionally, IF for Oct6 (UL marker, Dominguez., et al., 2013) and Sox5 (DL marker, Kwan et al., 2008) further strengthened our findings (Fig. S2A). These data indicate that Neurod2/6 deficiency causes expansion of UL neurons at expense of DL neurons during early cortical development.

**Figure 2.**
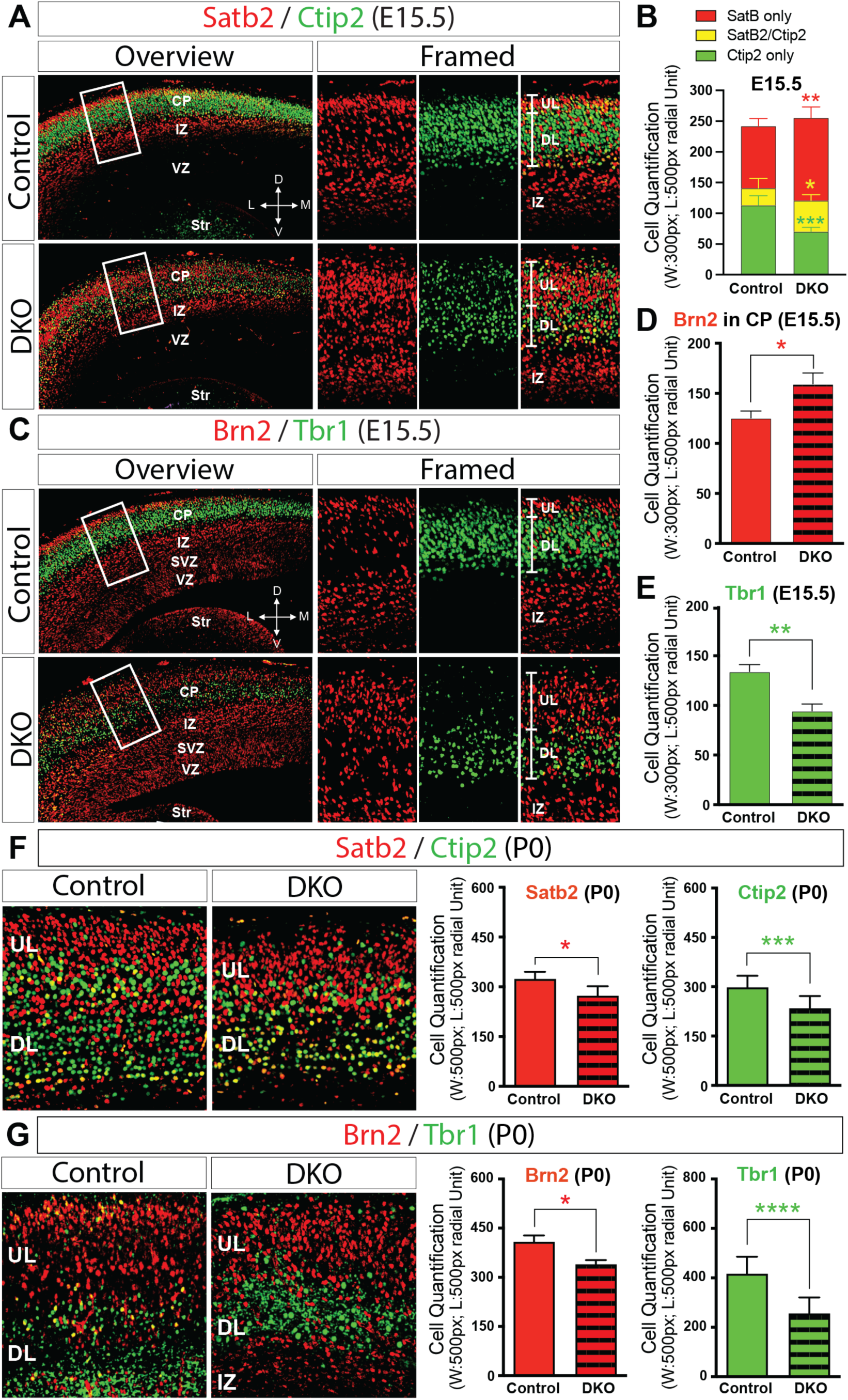
Cortical cell fate is misprogrammed in Neurod2/6 double deficient mice. **(A, B)** Immunofluorescence staining (IF, A) for Satb2 and Ctip2 on coronal sections of control and Neurod2/6 double deficient (DKO) brains shows increased Satb2+ upper layer neurons (UL, red) and decreased Ctip2+ deeper layer neurons (DL, green) in DKO brains at E15.5. (B) Cell quantification shows the number of single Satb2+ UL cells in DKO brains (134.90 ± 7.93, n = 5) is increased by 25.11% ± 7.18% (*p* = 0.0081, **) relative to that of control brains (101.00 ± 5.56, n = 5), nevertheless, the number of single Ctip2+ DL cells in DKO brains (69.80 ± 3.25, n = 5) is decreased by 38.20% ± 6.89% (p = 0.00050, ***) relative to that of control brains (112.90 ± 7.07, n = 5). Moreover, the number of Satb2/Ctip2 double positive cells in DKO brains (50.80 ± 4.34) is also increased by 81.43% ± 29.81% (*p* = 0.026, *) in comparison to that of control brains (28.00 ± 7.13). **CP**, cortical plate; **IZ**, intermediate zone; **VZ**, ventricular zone; **Str**, striatum. **D**, dorsal; **L**, lateral; **V**, ventral; **M**, medial. (**C-E**) IF for Brn2 and Tbr1 at E15.5 also shows increased Brn2+ UL cells (red) and decreased Tbr1+ DL cells (green) in DKO brains. The number of Brn2+ UL cells in DKO brains (159.50 ± 10.96, n = 5) is increased by 27.17% ± 10.35% (*p* = 0.030, *) relative to that of control brains (125.40 ± 6.94, n = 5), while the number of Tbr1+ DL cells in DKO brains (94.47 ± 15.53, n = 5) is decreased by 29.50% ± 7.39% (*p* = 0.0040, **) relative to that of control brains (134.00 ± 15.78, n = 5). **SVZ**, subventricular zone. **(F, G)** IF for Satb2/Ctip2 (F) and Brn2/Tbr1 (G) on coronal sections of control and DKO brains shows both UL and DL cells are reduced in DKO brains at P0. The Satb2+ UL and Ctip2+ DL cells are decreased by 15.61% ± 11.07% (*p* = 0.041, *; n = 8) and 21.31% ± 17.17% (*p* = 0.00080, ***; n = 8), respectively. Moreover, the Brn2+ UL and Tbr1+ DL cells are decreased by 16.82% ± 5.78% (*p* = 0.011, *; n = 6) and 38.68% ± 22.64% (*p* < 0.0001, ****; n = 6), respectively.

However, at later stages of development not only the number of DL neurons was reduced relative to the controls, but also does the number of UL neurons. IF for Satb2 and Ctip2 demonstrated that UL and DL neurons were both reduced in DKO brains perinatally. Satb2+ UL and Ctip2+ DL neurons were decreased by 15.61% ± 11.07% (*p* = 0.041, *) and by 21.31% ± 17.17% (*p* = 0.00080, ***) in DKO brains, respectively, compared with controls (Fig. 2F). Likewise, Brn2+ UL and Tbr1+ DL cells were decreased by 16.82% ± 5.78% (*p* = 0.011, *) and by 38.68% ± 22.64% (*p* < 0.0001, ****), respectively, in DKO brains at P0 (Fig. 2G). IF for other layer markers, such as Oct6, Foxp2 (DL marker, Kast et al., 2019) and Sox5, confirmed these findings (Fig. S2B, S2C). Our data indicate that Neurod2/6 control the specification of neocortical subtypes of projection neurons.

### Neurod2/6 deficiency does not affect Pax6+ apical progenitors

Cortical laminar subtype specification is coordinated with their molecular expression profile, which has been largely programmed in the dividing progenitors (Molyneaux et al., 2007; Telley et al., 2016; Oberst et al., 2019). Hence we investigated if APs are also affected in the neocortex of Neurod2/6 DKO mice. The expression of Pax6, the AP marker and cell identity determinant in the VZ (Götz et al., 1998), was not changed in DKO cortices as compared to the control brains throughout embryonic neurogenesis (Fig. S3A). Another landmark of cell fate specification is an expression gradient of phosphorylated Erk1/2 (pErk1/2) in the developing neocortex. It is essential for neuronal differentiation of APs and laminar specification (Ortega et al., 2010; Xing et al., 2016; Gantner et al., 2021; Parthasarathy et al., 2021). The graded expression of pErk1/2 was not altered in DKO brains (Fig. S3B). In addition, we analyzed if Neurod2/6 regulate the expression of genes that are involved in the control of cortical area specification in the VZ (O’Leary et al., 2007; Caronia-Brown et al., 2014). Neither lateral high to medial low gradients of expression of area regulators, such as Coup-TF and Pax6 (Fig. S4A-A’ and S4B-B’), nor medial high to lateral low graded regulators, such as Etv5 and Emx1 (Fig. S4C-C’ and S4D-D’), were changed in DKO cortices as compared to the control brains. Taken together, our data indicate that Neurod2/6 are not involved in direct regulation of APs.

### Neurod2/6 regulate basal progenitor differentiation and laminar fate specification cell-autonomously

The NeuroD members are key TFs that facilitate neuronal maturation and axonal navigation cell-autonomously (Tutukova et al., 2021; Pieper et al., 2019; Yan et al., 2023). Therefore, we asked if Neurod2/6 also regulate BP differentiation and cell fate specification in a cell-autonomous manner. In order to address this, we first analyzed the expression of Neurod2 or Cre (that recapitulates Neurod6 expression in NeuroD6^Cre^ mouse line) in APs and BPs. Both Neurod2 and Cre display mutually exclusive expression patterns with Pax6 (Fig. 3A, 3C). However, Neurod2 and Cre have been found to colocalize with Tbr2 in a subpopulation of BPs throughout neurogenesis at the border between BPs and newborn neurons starting from E11.5 (Fig. 3B, 3D). Notably, the cells co-expressing Cre/Tbr2 are progressively increasing in Neurod2/6 DKO cortex, implicating that Tbr2+ ectopic cells in DKO brains are indeed those that fail to differentiate and migrate due to intrinsic deficiency of Neurod2/6 (Fig. 3D). Extrinsic factors, such as radial glia fibers and Reelin, in cerebral cortex are known to be essential for proper positioning of newborn neurons (Tissir et al., *review*, 2003; Rakic, *review*, 2009; Belvindrah et al., 2007; Liu et al., 2015). In order to test if these can be changed in DKO brains, we performed IF for – Nestin, Blbp (brain lipid binding protein) (Kriegstein et al., 2003; Anthony et al., 2005), and Reelin. We found radial glia scaffold as well as Reelin expression to be normal throughout corticogenesis in DKO brains (Fig. S5).

**Figure 3.**
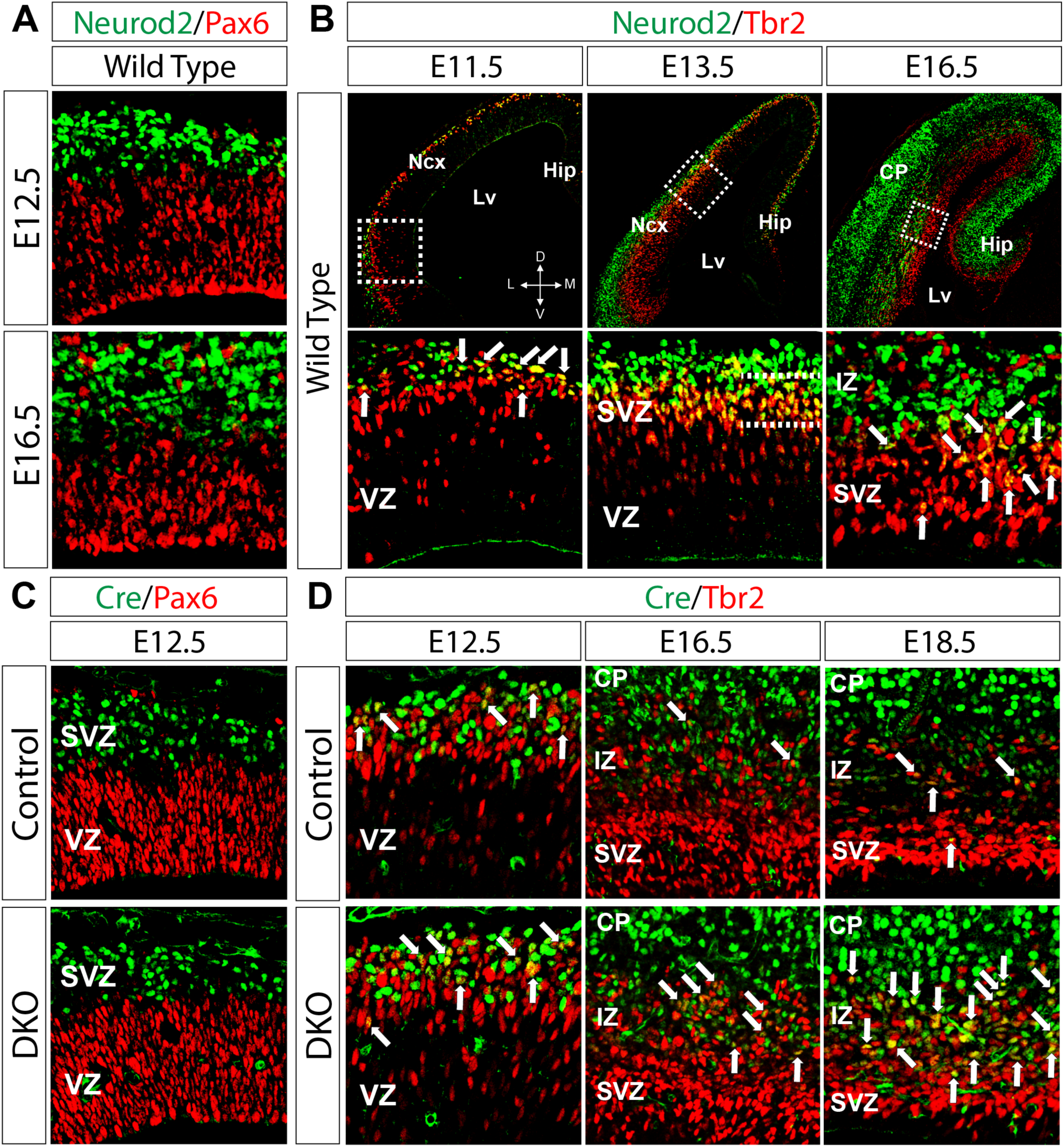
Neurod2/6 are transiently expressed in Tbr2+ basal progenitors. **(A)** IF for Neurod2 (green) and Pax6 (red) on wild type brain sections at E12.5 and E16.5 shows that Neurod2 is barely expressed in Pax6+ apical progenitors (APs). **(B)** IF for Neurod2 and Tbr2 (red) on wild type brain sections at E11.5, E13.5 and E16.5 shows that a subpopulation of Tbr2+ BPs in the SVZ during corticogenesis also express Neurod2. The images in lower panel represent the magnified views of framed parts in upper panel. **(C)** IF for Cre (Neurod6, green) and Pax6 on coronal sections of control and DKO brains at E12.5 shows that Neurod6 is never present in Pax6+ APs. **(D)** IF for Cre and Tbr2 on control and DKO brain sections at E12.5, E16.5 and E18.5 shows that Neurod6 is transiently expressed in Tbr2+ BPs throughout cortical neurogenesis in control brains (upper panel), and that Cre/Tbr2 double positive cells are accumulating during development in DKO brains (lower panel).

We next performed *in utero* electroporation (IUE) with the constructs that enable Cre-dependent expression of either GFP or bicistronic Neurod6-GFP at E12.5 and analyzed the brains at E15.5. While IUE of GFP into control and DKO brains did not disturb the expression of Tbr2 (Fig. 4A) or Pax6 (Fig. S6A), enforced Neurod6 expression caused depletion of Tbr2+ BPs (Fig. 4B), but not Pax6+ APs (Fig. S6B). In line with this, ISH for Tbr2 on Neurod6 electroporated brains displayed downregulation of Tbr2 expression at electroporation sites, but not in the neighboring areas (Fig. S6C). In order to test if Neurod6 restoration in DKO brains can rescue the defects of cell fate specification, we performed co-IF for GFP and laminar markers on the electroporated brains. We found that the proportion of GFP+/Satb2+ cells in GFP-electroporated DKO brains (∼77.90%) was much higher than that in GFP-electroporated control brains (∼52.11%), whereas the proportion of GFP+/Ctip2+ cells in GFP-electroporated DKO brains (∼46.44%) was lower than that in control brains (∼56.62%), mimicking the similar deficits of laminar fate decision in DKO brains as in Fig 2A – 2E. IUE of Neurod6 into DKO brains reinstated the normal proportions of Satb2+ or Ctip2+ cells (GFP+/Satb2+: ∼56.65% and GFP+/Ctip2+: ∼59.32%) comparable to those in GFP-electroporated controls (Fig. 4C-4E). Moreover, the rescue effects of Neurod6 were further underpinned by the evidence on additional laminar markers, such as Brn2 and Tbr1 (Fig. 4F-4H). Such acute and focal Neurod6 restoration in a limited subpopulation of progenitors could rescue the cellular deficits in cell fate specification, indicating that Neurod2/6 control cell fate cell-autonomously. Our data also indicate that Neurod2/6 are critical for sequential production of subtypes of neocortical projection neurons.

**Figure 4.**
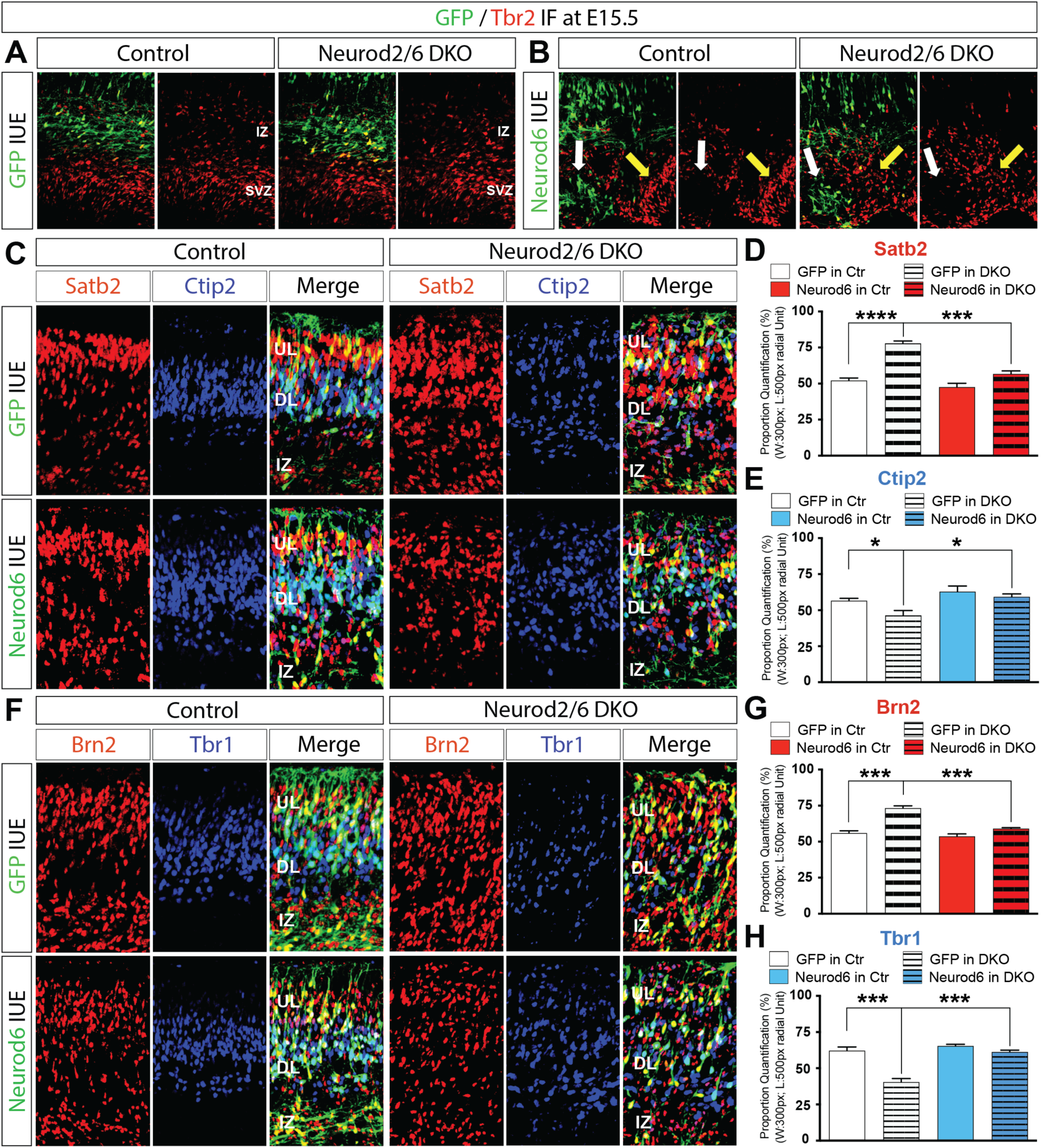
Restoration of Neurod6 expression in DKO cortices at an earlier neurogenic stage rescues cell specification deficits. **(A, B)** GFP or bicistronic Neurod6-GFP was electroporated into control and DKO brains at E12.5, which were then sampled at E15.5 (the same in **C - H**). IF for GFP and Tbr2 on GFP- or Neurod6-electroporated brain sections shows that Neurod6 over-expression in both control and DKO brains causes depletion of Tbr2+ BPs at IUE sites (white arrowheads) but not in neighboring non-IUE regions (yellow arrowheads, **B**), while GFP IUE does not disturb the distribution of Tbr2+ BPs (**A**). **(C-E)** IF for GFP, Satb2 and Ctip2 on GFP- or Neurod6-electroporated brain sections (at E15.5) shows in utero electroporation (IUE) of Neurod6 in DKO brains downregulates UL neurons but upregulates DL neurons (**C**). Proportional quantification of Satb2/GFP (**D**) or Ctip2/GFP (**E**) double positive cells relative to all GFP+ cells shows the ratio of Satb2/GFP+ UL cells in GFP-electroporated DKO brains is 77.90% ± 3.22% (n = 4) in comparison to GFP-electroporated control brains (52.11% ± 3.47%, n = 4), but Neurod6 IUE reduced the ratio to 56.65% ± 4.11% (*p* = 0.00019, ***; n = 4). On the other hand, the ratio of Ctip2/GFP+ DL cells in GFP-electroporated DKO brains is 46.44% ± 6.67% (n = 4) in comparison to GFP-electroporated control brains (56.62% ± 3.21%, n = 4), whereas the ratio is elevated to 59.32% ± 3.92% (*p* = 0.016, *; n = 4) when Neurod6 was electroporated in DKO brains. **(F-H)** IF for GFP, Brn2 and Tbr1 on GFP- or Neurod6-electroporated brain sections confirms the rescue effects of Neurod6 IUE in DKO brains (**F**). The ratio of Brn2/GFP+ UL cells in GFP-electroporated DKO brains is 73.29% ± 3.11% (n = 4) in comparison to GFP-electroporated control brains (55.97% ± 3.12%, n = 4), however, Neurod6 IUE reduced the ratio to 59.03% ± 1.40% (*p* = 0.00020, ***; n = 4) (**G**). The ratio of Tbr1/GFP+ DL cells in GFP-electroporated DKO brains is 40.45% ± 4.89% (n = 4) in comparison to GFP-electroporated control brains (62.18% ± 5.02%, n = 4), but Neurod6 restoration in DKO brains elevated the ratio up to 61.33% ± 2.40% (*p* = 0.00029, ***; n = 4) (**H**).

### Neurod2/6 promote timely specification of basal progenitors into upper layer neurons

Though Tbr2+ BPs give rise to neurons in all cortical layers, they mainly produce UL neurons after E14.5 through neurogenic divisions (Sessa et al., 2008; Kowalczyk et al., 2009; Mihalas et al., 2016). However, a large population of BPs in Neurod2/6 DKO cortices retain their Tbr2 expression and are arrested in SVZ/IZ (Fig. 1). We thus reasoned if UL neuron reduction in DKO brains (Fig 2F, 2G) is resulted from failure of Tbr2+ BP differentiation. Tuj1, the earliest neuronal fate marker (Memberg et al., 1995; Reyes et al., 2008), is strongly expressed in postmitotic compartment but minimally in Tbr2+ BPs (Fig. 5A). Enforced Neurod2 expression in both control and DKO brains led to precocious activation of Tuj1 expression in Tbr2+ BPs at the IUE sites, but not those in the neighboring cells (Fig. 5B, 5C), indicating Neurod2/6 expression is sufficient to drive BPs into neurogenic differentiation. To further find out if the ectopic BPs born after E14.5 are properly specified to UL neurons in the presence of Neurod2, we performed IF for GFP and Satb2 to analyze the brains that were electroporated with GFP or bicistronic Neurod2-GFP at E14.5 and collected at E18.5 (Fig. 5D). The proportion of GFP+/Satb2+ cells was ∼77.48% in GFP-electroporated control brains but significantly decreased to ∼35.18% in Neurod2/6 DKO brains (*p* < 0.0001, ****), largely due to the presence of Satb2 fated cells arrested in the SVZ/IZ. Notably, Neurod2 restoration in DKO brains increased the proportion of GFP+/Satb2+ cells up to the similar level (∼75.11%) with that in GFP-electroporated control brains (Fig. 5E).

**Figure 5.**
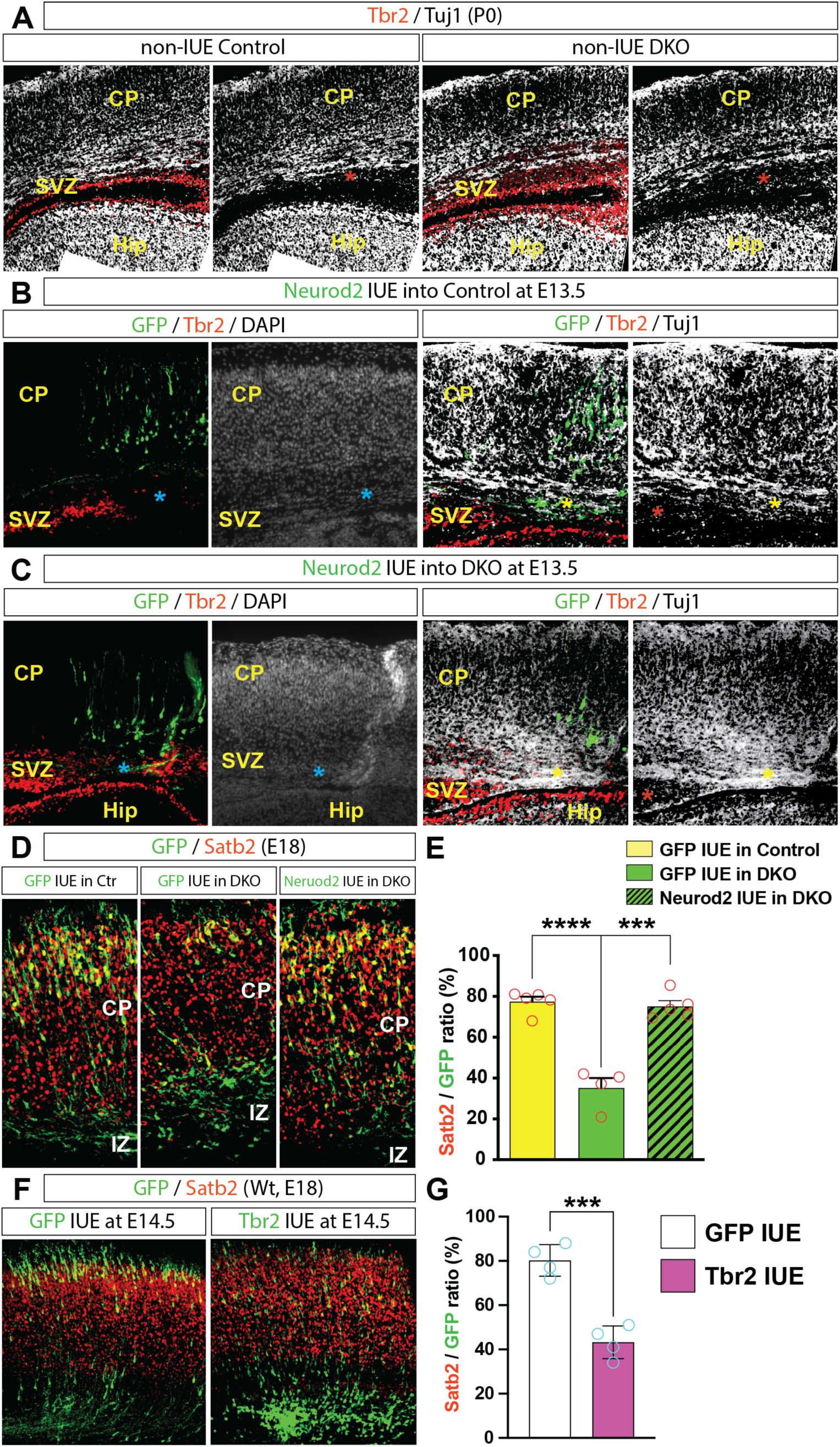
Electroporated Neurod2 at a later neurogenic stage instructs basal progenitor differentiation into upper layer neurons. **(A)** IF for Tbr2 and Tuj1 on control and DKO brain sections at P0 shows that Tuj1 expression is extremely weak in the SVZ enriched with Tbr2+ BPs (red asterisks). **(B, C)** Bicistronic Neurod2-GFP was electroporated into control (**B**) and DKO (**C**) brains at E13.5, which were sampled at E18.5 (the same in **D, E**). IF for GFP, Tbr2 and Tuj1 on Neurod2-electroporated brain sections shows that IUE of Neurod2 in both control and DKO brains results in onset of Tuj1 expression in Tbr2+ BPs at IUE sites (blue asterisks) but not in adjacent SVZ. DAPI nuclei counterstaining on the neighboring sections shows that there are cell populations at IUE sites to avoid an artifact caused by IUE. These cells at IUE sites instead express Tuj1, an earliest neurogenic marker (yellow asterisks), but not Tbr2, in contrast to neighboring Tbr2+ cells in SVZ (red asterisks). **(D, E)** GFP and bicistronic Neurod2-GFP was electroporated into control and DKO brains at E14.5, which were then sampled at E18.5. IF for GFP and Satb2 shows that the proportion of GFP+/Satb2+ relative to all GFP-electroporated cells in DKO brains (∼35.18%, n = 4) is significantly reduced in comparison to that in control brains (∼77.48%, n = 5) (*p* < 0.0001, ****). IUE of Neurod2 into DKO brains promotes Satb2+ cell differentiation and increases the ratio up to ∼75.11% (*p* = 0.00010, ***; n = 5), similar to control brains. **(F, G)** GFP and bicistronic Tbr2-GFP was electroporated into wild type brains at E14.5, which were then sampled at E18.5. IF for GFP and Satb2 shows that Tbr2 IUE restricts the majority of cells beneath CP, resulting in reduced ratio of Satb2+ UL cells by 46.11% ± 6.41% (*p* = 0.00040, ***; n = 4) relative to total GFP+ cells in comparison to that of GFP-electroporated controls.

On the other hand, we studied if excessive Tbr2 expression in BPs affects UL neuron commitment. We electroporated GFP or bicistronic Tbr2-GFP in wild type brains at E14.5 and analyzed the brains at E18.5 (Fig. 5F). Tbr2 over-expression *in vivo* prevented radial migration of cortical projection neurons and impaired their neurite arborization (Fig. S7), highly consistent with the previous report (Sessa, et al., 2017). Importantly, IUE of bicistronic Tbr2-GFP retained a large population of GFP+/Satb2-cells in the SVZ/IZ and thus led to significantly reduced production of Satb2+ UL cells (by 46.11% ± 6.41%; *p* = 0.00040, ***) in comparison to GFP-electroporated brains (Fig. 5G), resembling the deficits of UL projection neuron specification in Neuod2/6 DKO brains. Taken together, Neurod2/6 are essential for timely specification of Tbr2 BPs into UL neurons at later neurogenic stages of neocortex.

### Olig2+ glial precursors are ectopically upregulated in the cortex of Neurod2/6 deficient mice

Next we reasoned if the ectopic Tbr2+ cells that fail to differentiate into neurons in Neurod2/6 deficient brains could retain their proliferating activity. We performed co-IF for Tbr2 and proliferation markers, such as PCNA (proliferating cell nuclear antigen) or Ki67 (Gerdes et al., 1991; Maga et al., 2003; Juríková et al., 2016; Sobecki et al., 2016). We detected substantial numbers of Tbr2+ cells, though not all, expressing either PCNA and/or Ki67 in the IZ of DKO brains at P0 (Fig. 6A, 6B). Sox2 is a key regulator for self-renewal and proliferation of neural progenitors (Taranova et al., 2006; Hagey et al., 2014; Pagin et al., 2021). IF for Tbr2 and Sox2 detected increase in number of Tbr2+/Sox2+ cells in the IZ of DKO brains as compared with controls (Fig. 6C). Taken together, our data indicate that ectopic Tbr2+ cells in DKO brains, at least in part, retain their proliferating activity.

**Figure 6.**
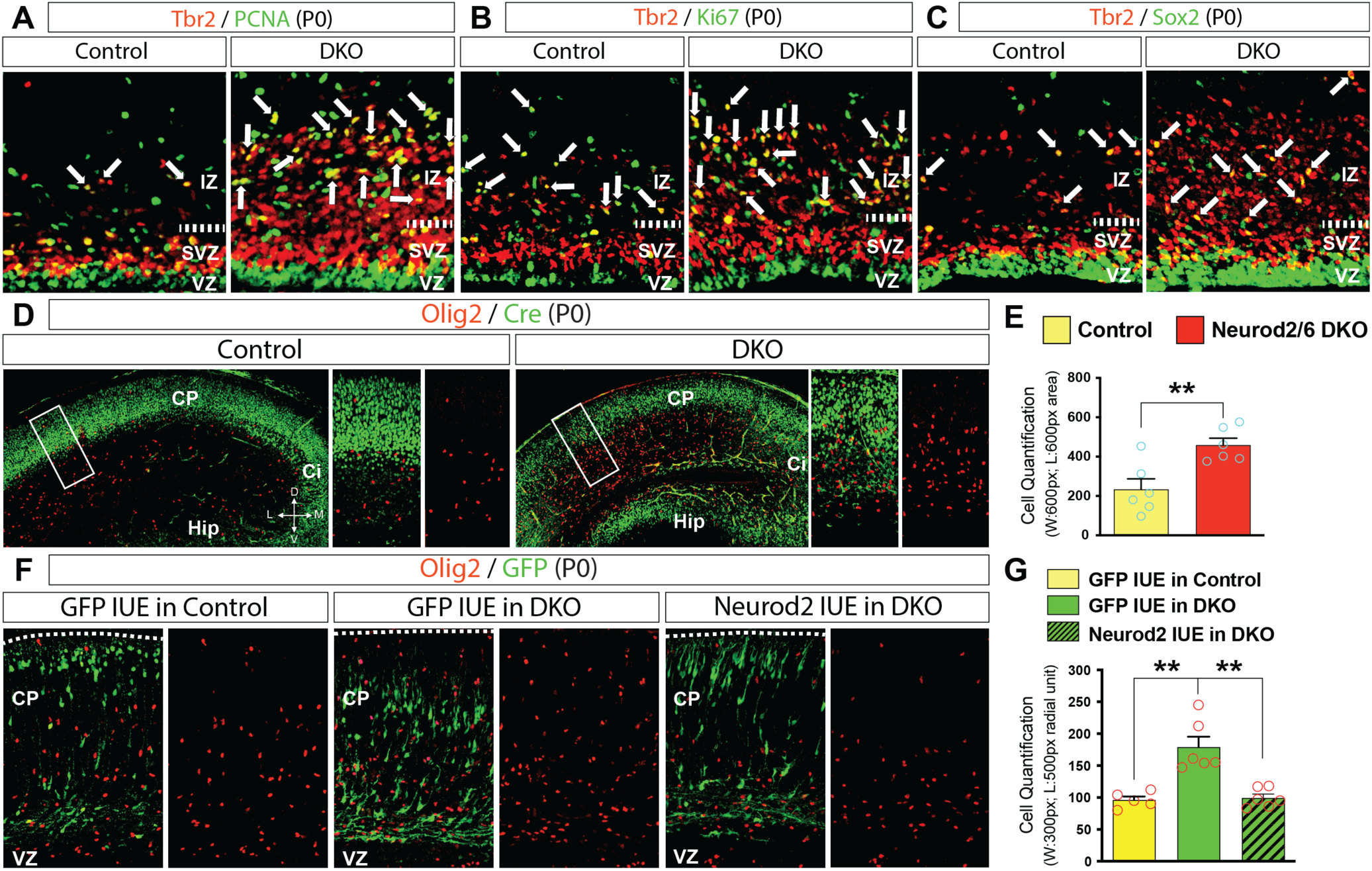
Olig2+ glial progenitors are ectopically elevated in Neurod2/6 DKO brains. **(A-C)** Co-IF for Tbr2/PCNA (**A**), Tbr2/Ki67 (**B**) and Tbr2/Sox2 (**C**) on control and DKO cerebral coronal sections at E18.5 shows that a large subpopulation of ectopic Tbr2+ cells co-express PCNA, Ki67 and Sox2 in the IZ of DKO brains. Each arrowhead indicates one or one cluster of double positive cells in the IZ. **(D, E)** IF for Cre (green) and Olig2 (red) on control and DKO cerebral coronal sections at P0 (**D**). (**E**) Quantification of Olig2+ glial progenitors for control and DKO cortices at P0. The number of Olig2+ cells in DKO brains is upregulated to 458.80 ± 34.89 (*p* = 0.0053, **; n = 6) in contrast to that of controls (233.80 ± 52.92, n = 6). **(F, G)** GFP or bicistronic Neurod2-GFP was electroporated into control and DKO brains at E14.5, which were later sampled at P0. IF for GFP (green) and Olig2 (red) on GFP- or Neurod2-electroporated brain sections shows the Olig2+ cells in GFP-electroporated region of DKO brains are upregulated in comparison to GFP-electroporated control brains, which is normalized by Neurod2 IUE in DKO brains (**F**). (**G**) Quantification of Olig2+ cells in the GFP+ regions of GFP or Neurod2-GFP electroporated control and DKO brains shows that the number of Olig2+ cells in GFP-electroporated DKO brain areas is 179.00 ± 39.98 (*p* = 0.0017, **; n = 6) in contrast to that of GFP-electroporated control brain areas (96.20 ± 12.38, n = 5), however, the number in Neurod2-electroporated DKO brain areas is reduced to 99.33 ± 15.03 (*p* = 0.0010, **; n = 6).

Delay in the mitotic cycle exit or ectopic progenitor proliferation often causes expansion of cell numbers of later born subtypes (Nieto et al., 2001; Miller et al., 2007; Kohwi et al., 2013). Considering BPs can produce glial lineage following UL neurogenesis (Li, et al., 2021), we therefore hypothesized that more proliferative BPs in DKO brains could over-produce glial cells at the late stage of embryonic corticogenesis. To verify this, we performed IF for Cre and Olig2 - a marker for glial precursors and oligodendrocytes (Marshall et al., 2005; Ono et al., 2009; Seuntjens et al., 2009).

The number of Olig2+ cells in DKO brains was almost doubled of that in control brains at P0 (*p* = 0.0053, **; Fig. 6D, 6E). Notably, Cre expression (Neurod6-expressing excitatory neurons) never overlapped with Olig2 expression, indicating that Neurod2/6 do not act in Olig2+ cells directly to regulate their cell fate or production. On the other hand, IUE of Neurod6 into the progenitors in DKO brains downregulated the number of Olig2+ cells in the electroporation area to the similar level as that in the GFP-electroporated controls (Fig. 6F, 6G). These data indicate that Neurod2/6-driven genetic program suppresses glial cell fate in BPs and controls a proper timely switch from neurogenesis to gliogenesis.

## Discussion

Neurod2/6 expression in cerebral cortex begins with the onset of neurogenesis in the SVZ and retains in the postmitotic neurons (Wu et al., 2005; Goebbels et al., 2006; Bormuth et al., 2013).

These TFs have been long known to regulate multiple cellular processes in corticogenesis, such as neuronal differentiation, migration, axon navigation and synapse plasticity (Wu et al., 2005; Ince-Dunn et al., 2006; Bormuth et al., 2013; Guzelsoy et al., 2019; Yan et al., 2023). However, their downstream mechanisms underlying proneural differentiation and their potential role in sequential specification of cortical cell subtypes are poorly studied. One of the aims of this study was to test the hypothesis that Neurod2/6 might control differentiation of specific subtypes of neural progenitors. Indeed we could show that Neurod2/6 do not have a direct role in differentiation of APs, but specifically regulate BP differentiation and thus the cell fates of BPs’ progeny.

BPs are derived from Pax6+ APs and populated in the SVZ. They may either produce neurons in one neurogenic division or undergo one or more self-renewing divisions. BPs thus serve as a secondary progenitor pool to amplify neuron production (Noctor et al., 2004; Sessa et al., 2017). How-ever, how a BP cell makes the neurogenic versus proliferative decision is unclear. Our mouse model turned out to be a unique tool to study the mechanisms and effects of endogenous BP expansion.

Tbr2 is a signature gene for BPs, and has been reported to be a key TF that facilitates BP genesis and self-renewal, supported by the findings that Tbr2 deletion in forebrain resulted in reduced BP numbers and mitoses, and consequently reduced number of neurons in all cortical layers (Arnold et al., 2008; Sessa et al., 2008). Furthermore, Tbr2 over-expression by IUE increased mitoses in SVZ and inhibited the genes accelerating cell cycle exit, such as Btg2 and Cdkn1b (Arnold et al., 2008; Sessa et al., 2008 and 2017). Consistently, we found that ectopic BPs in DKO brains retained Tbr2 expression and failed to differentiate. These cells instead reentered proliferative cycling. On the other hand, Neurod2/6 were expressed in a subset of BPs throughout cortical neurogenesis and promoted BP cell cycle exit. In addition, acute Neurod2/6 over-expression downregulated Tbr2 expression and converted almost all Neurod2/6+ BPs into neurons. Collectively, these results indicate that Neurod2/6 trigger the immediate neurogenic process in BPs by suppressing Tbr2-dependent proliferative program.

It has been shown that BPs produce cortical neurons of all layers with progressive cell fate restriction (Noctor et al., 2004; Mihalas et al., 2016; Sessa et al., 2017). Early born BPs (E11.5 – E13.5) may be specified either into DL neurons in one round rapid division or into UL neurons due to self-renewal or protracted differentiation, but late BPs (since E15.5) exclusively produce UL neurons (Mihalas et al., 2016). The same report also indicated that Tbr2 critically regulated this tempo of BP neurogenesis and thus the laminar fates of daughter cells. One of their main findings to support this idea was that Tbr2 deficiency in BPs caused expanded DL neuron production (birthday mainly at E12.5) at the expense of UL neurons. Intriguingly, we showed that Tbr2 upregulation due to Neurod2/6 deficiency exerted an opposite effect on laminar fates to Tbr2 deficiency, and that the deficits in laminar fate choice could be almost completely rescued by Neurod2/6 restoration at E12.5. These data suggest that Neurod2/6 regulate timely acquisition of cortical DL subtypes upstream of Tbr2 during early neurogenesis.

The reduced UL neurons in Tbr2 deficient mouse brains were resulted either from inadequate BP amplification (Arnold et al., 2008; Sessa et al., 2008) or from impaired timely differentiation (Mihalas et al., 2016) during early neurogenesis (E11.5 – E13.5), but Tbr2’s roles in late neurogenesis (E15.5 – E17.5) is not thoroughly elucidated. Here we demonstrated Tbr2 IUE at E14.5 prevented progenitors from differentiation into Satb2+ UL neurons. Additionally, others and we showed that Tbr2 electroporation led to impaired radial migration and neurite outgrowth of cortical pyramidal neurons (Sessa et al., 2017 and Fig. S7). These results suggest that Tbr2 over-expression in late neurogenesis may impede UL neuron differentiation and maturation. Notably, Tbr2+ BPs were arrested in SVZ/IZ and unable to differentiate into Satb2+ UL cells in Neurod2/6 DKO brains, but Neurod2/6 restoration at E14.5 completely rescued the defected generation of UL neurons. Neurod2/6 thus facilitate UL neurogenesis by driving BP differentiation after E14.5.

Glia cells arise from the progenitor cells in the VZ/SVZ following UL neuron generation during corticogenesis (Miller et al., 2007). The timing of switch from neurogenesis to gliogenesis in APs is critically regulated by proneural TFs, such as Ngn2 and Mash1. Ngn2/Mash1 double deficiency in APs caused premature and excessive astrogenesis at the expense of their neurogenic capacity (Nieto et al., 2001). BPs have been recently identified as a transient amplification source for Olig2+ glial precursors after E16.5 (Li et al., 2021; Liang et al., 2024). Here we show that deficiency of another group of proneural TFs (Neurod2/6) resulted in ectopic cycling of BPs and thus over-production of Olig2+ precursors. This finding is also supported by the IUE experiments that Neurod2/6 restoration in DKO brains reduced the local numbers of Tbr2+ BPs and Olig2+ glial precursors to a similar level with those of controls. Taken all together, Neurod2/6 govern sequential commitment of cortical subtype fates, such as DL versus UL neurons, neuronal versus glial, via regulating timely differentiation of Tbr2+ BPs.

## Notes

We here thank R Wunderlich, R Dannenberg, D Lajkó and J Schüller for excellent technical support. Neurod2-Null (Neurod2^NeoR/NeoR^) and Neurod6-Null (namely NEX-Cre, Neurod6^Cre/Cre^) mice were contributed by T Yonemasu and S Goebbels, respectively, and provided by KA Nave and MH Schwab, Max-Planck-Institute of Experimental Medicine, Göttingen, Germany. This study was financially supported by DFG (grant TA 303/14-1), and by the Russian Science Foundation, project № 21-65-00017 (VT and ST, genetic rescue experiments).

## Author contributions

**Kuo Yan:** 1) conceptualization of the project. 2) writing and editing of manuscript draft. 3) phenotypic characterization of Neurod2/6 mouse model. 4) IUE and investigation of IUE effects on Neurod2/6 mouse model. **Andrew G Newman:** 1) data analysis and interpretation **Svetlana Tutukova:** 1) Tbr2 IUE and experimental characterization of Tbr2 IUE effects on wt mice. **Maria Gavrish:** 1) Sample preparation **Ingo Bormuth and Olga Bormuth:** 1) Tbr2 ISH on Neurod2/6 mouse model. **Victor Tarabykin:** 1) conceptualization, supervision and administration of the project. 2) writing and editing of manuscript draft. 3) funding acquisition.

## Notes

### Competing Interest Statement

The authors have declared no competing interest.

### Summary of Updates

We revised the abstract, introduction, discussion of the main manuscript as well as the funding information.

