## Supplemental Materials for "Neurod2/6 transcription factors control basal progenitor differentiation and sequential production of neocortical cell subtypes"

### Supplementary Figures:

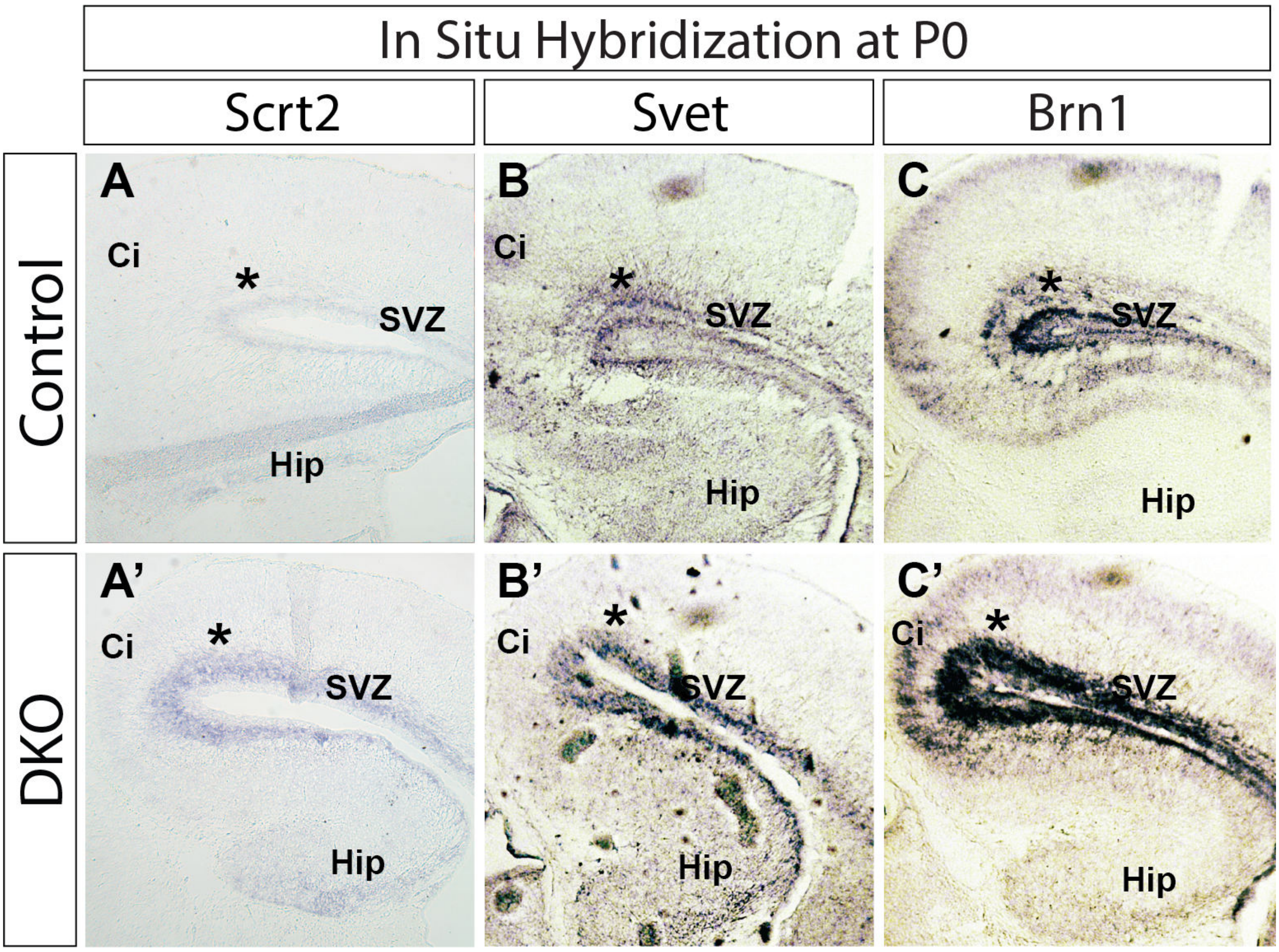


**Figure S1 Basal progenitor marker genes show expanded expression in Neurod2/6 double deficient brains.**

**(A – A' to C – C')** ISH shows three genes (Scrt2, Svet and Brn1) that are expressed in basal progenitors (BPs) are ectopically upregulated in Neurod2/6 double deficient (DKO) brains, consistent with the increased Tbr2+ cells in the subventricular zone (SVZ, asterisks) of DKO brains. **Hip**, hippocampus; **Ci**, cingulate cortex.

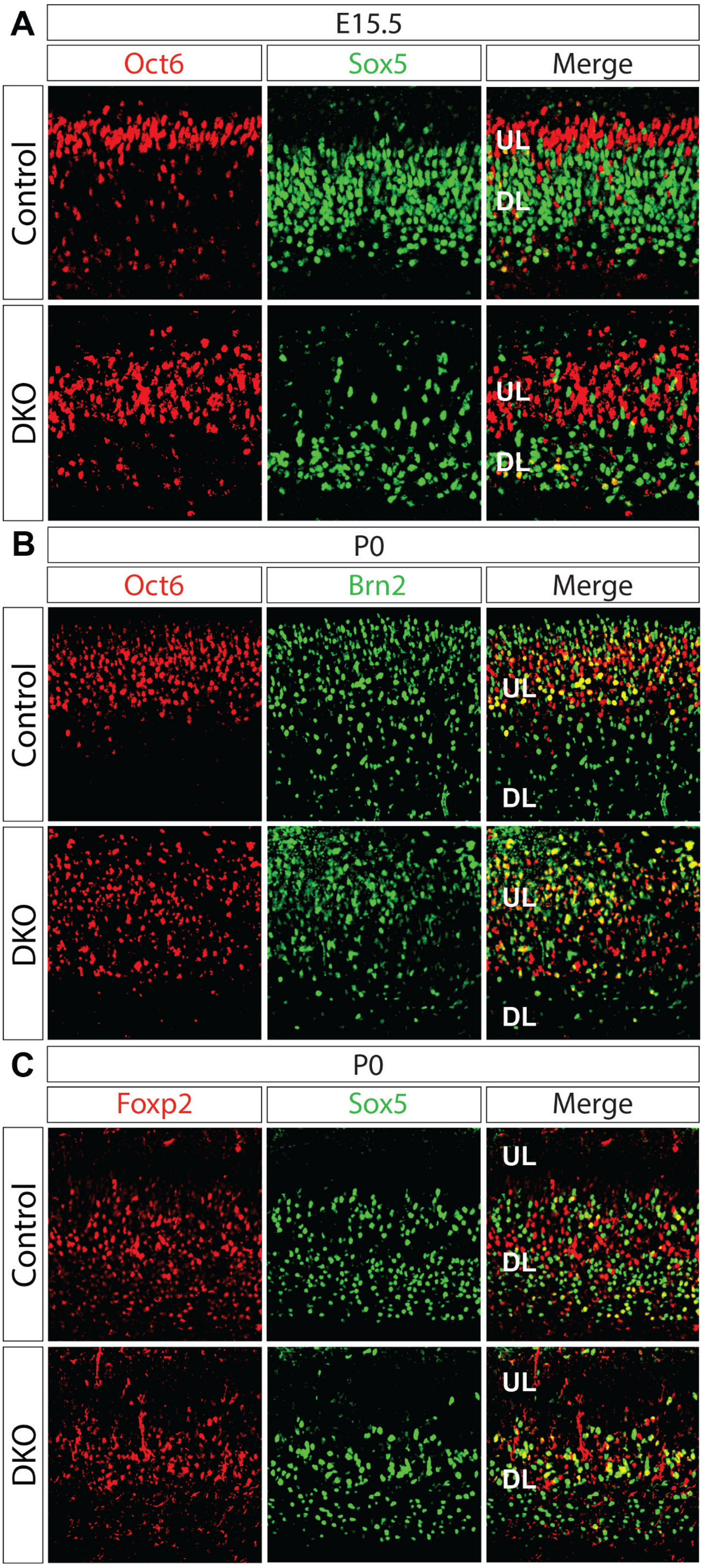
A
E15.5
Oct6
Sox5
Merge
Control
DKO
UL
DL
B
P0
Oct6
Brn2
Merge
Control
DKO
UL
DL
C
P0
Foxp2
Sox5
Merge
Control
DKO
UL
DL

10

**Figure S2 Cell fate specification is misregulated in Neurod2/6 double deficient brains.**

**(A)** Immunofluorescent staining (IF) for Oct6 and Sox5 on coronal sections of control and DKO brains shows increased Oct6+ upper layer cells (UL, red) and decreased Sox5+ deeper layer cells (DL, green) in DKO brains at E15.5.

**(B, C)** IF for Oct6/Brn2 (**B**) or Foxp2/Sox5 (**C**) on coronal sections of control and DKO brains shows both Oct6/Brn2-positive UL neurons and Foxp2/Sox5-positive DL neurons are visibly downregulated at P0.

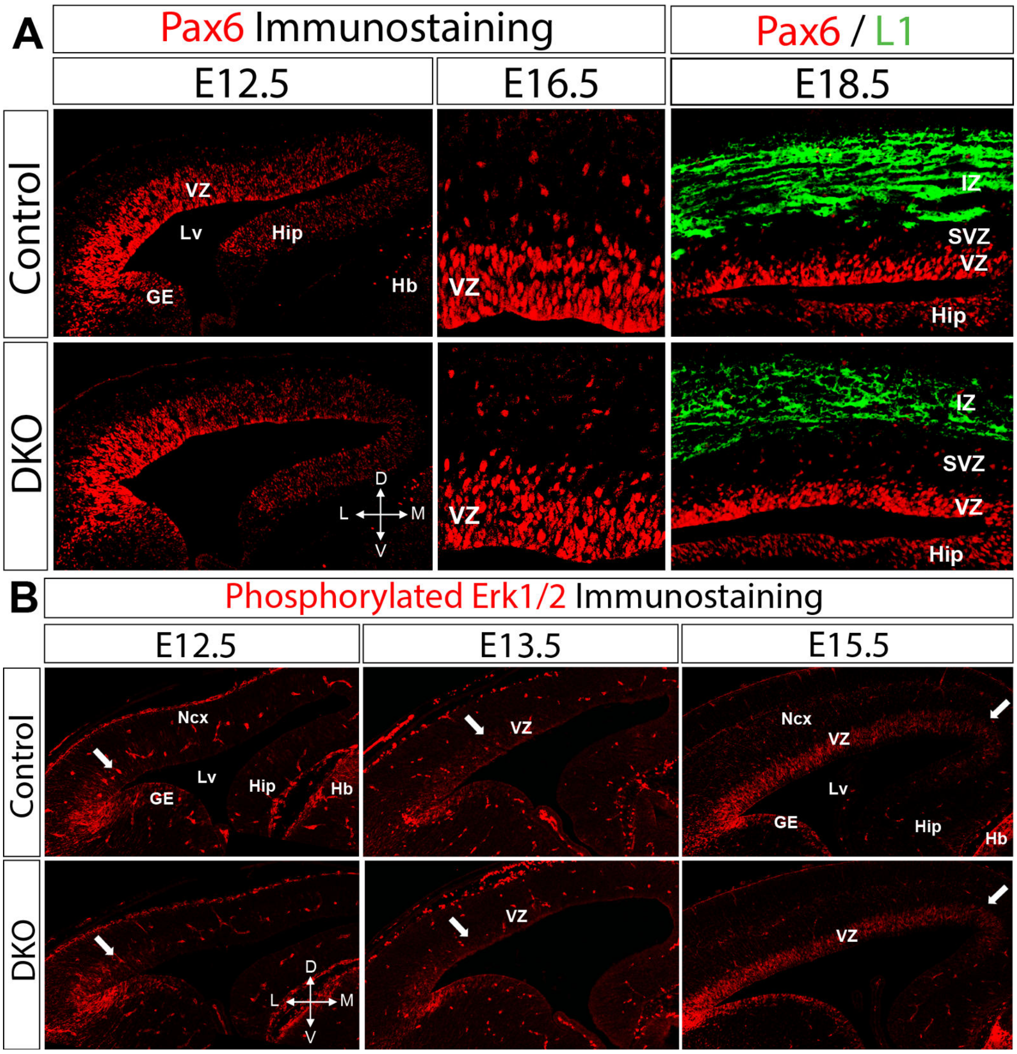


**Figure S3 Apical progenitors are normal in Neurod2/6 DKO brains.**

**(A)** IF for Pax6 (red) on cerebral coronal sections at E12.5, E16.5 and E18.5 shows that dynamic Pax6 expression in apical progenitors (APs) that are resident in ventricular zone (VZ) is unaltered in control and DKO mice. **Lv**, lateral ventricle; **GE**, ganglionic eminences; **Hb**, habenula; **IZ**, intermediate zone. **D**, dorsal; **M**, medial; **V**, ventral; **L**, lateral.

**(B)** IF for phosphorylated Erk1/2 (pErk1/2) on cerebral coronal sections at E12.5, E13.5 and E15.5 shows that the progressive expression of pErk1/2 in VZ is comparable in control and DKO mice during neurogenesis.

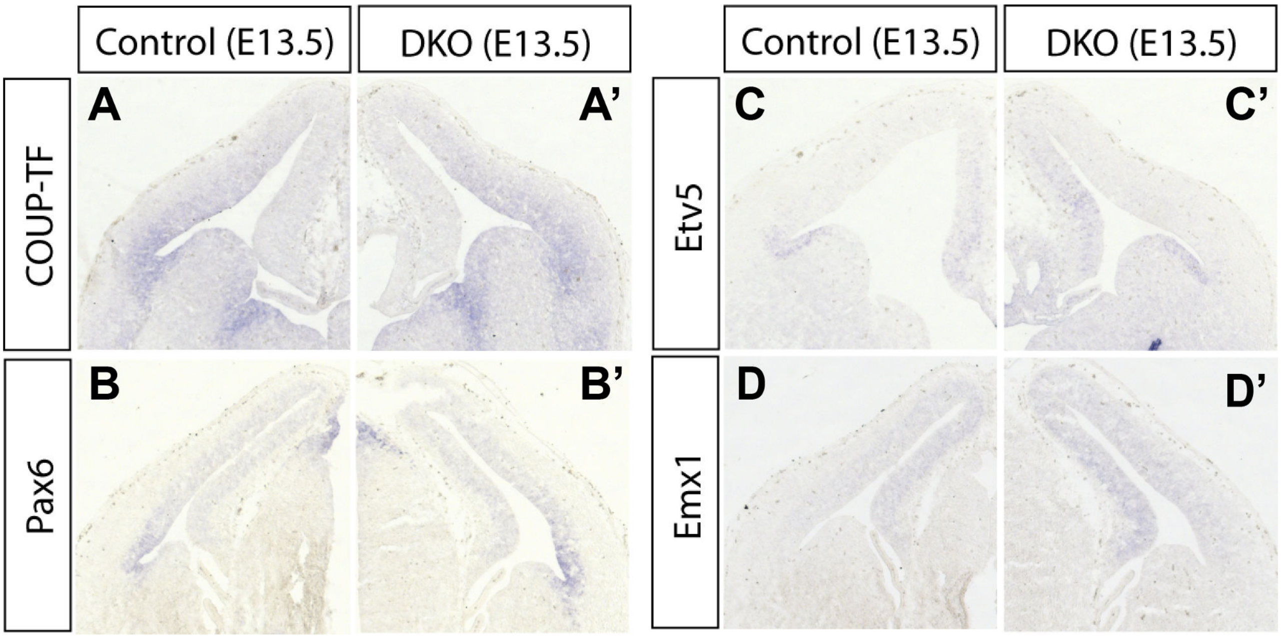


**Figure S4 The expression patterns of cortical regionalization regulators are normal.**

**(A – A’, B – B’)** *In situ* hybridization (ISH) shows that the lateral high to medial low graded expression of COUP-TF (**A – A’**) and Pax6 (**B – B’**) is comparable in control and Neurod2/6 DKO brains during early neurogenesis.

**(C – C’, D – D’)** The medial high to lateral low graded expression of Etv5 (**C – C’**) and Emx1 (**D – D’**) is also similar in Neurod2/6 DKO brains.

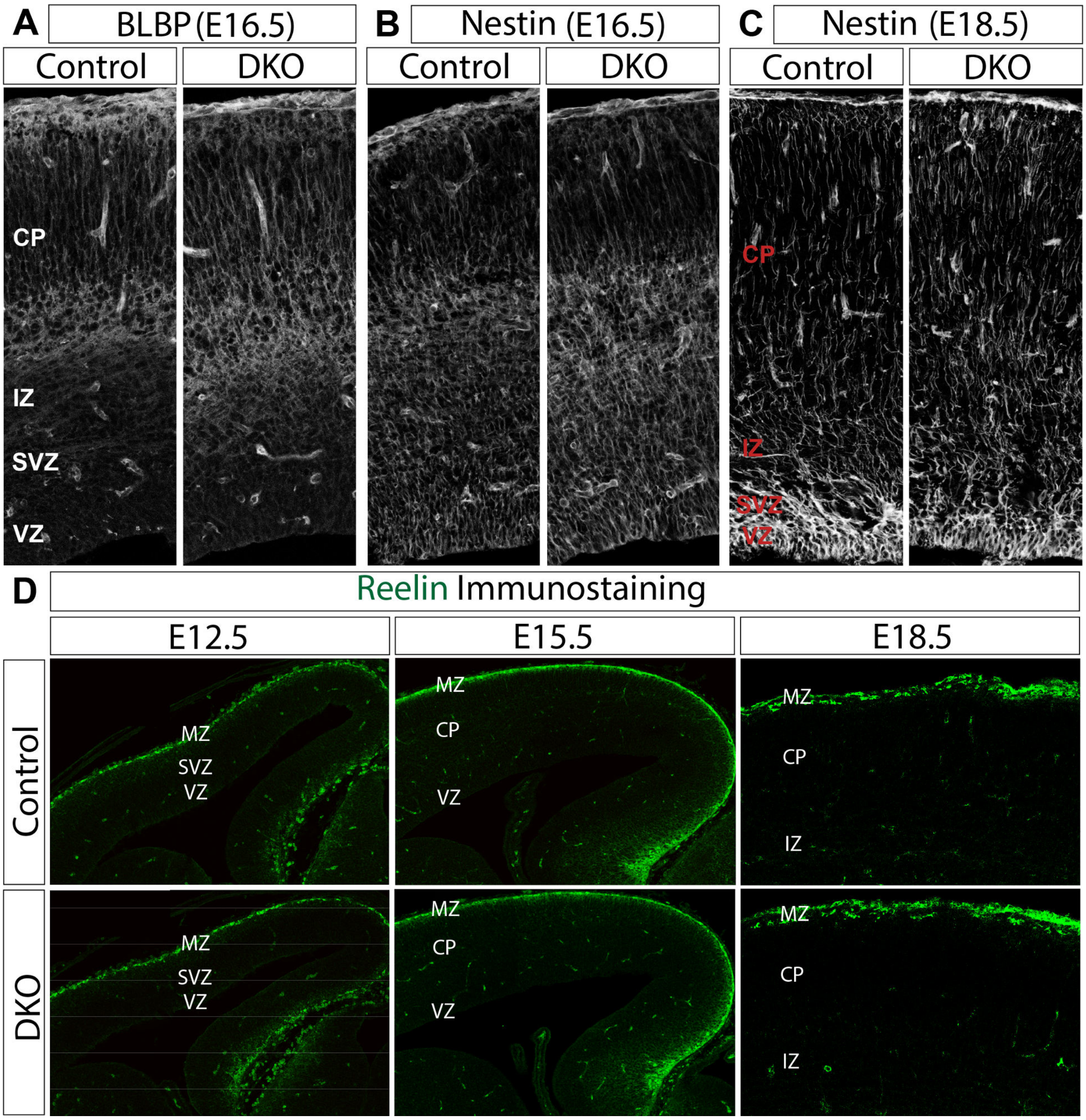


**Figure S5 Radial glia morphology and Reelin production are normal in Neurod2/6 DKO brains.**

**(A-C)** IF for BLBP at E16.5 (**A**), and Nestin at E16.5 (**B**) and E18.5 (**C**) on brain sections shows that the radial glial scaffold for cortical neuron migration are normal in DKO brains. CP, cortical plate.

**(D)** IF for Reelin on brain sections at E12.5, E15.5 and E18.5 shows that the production and distribution of Reelin+ Cajal-Retzius cells are also normal during corticogenesis in DKO brains. MZ, marginal zone.

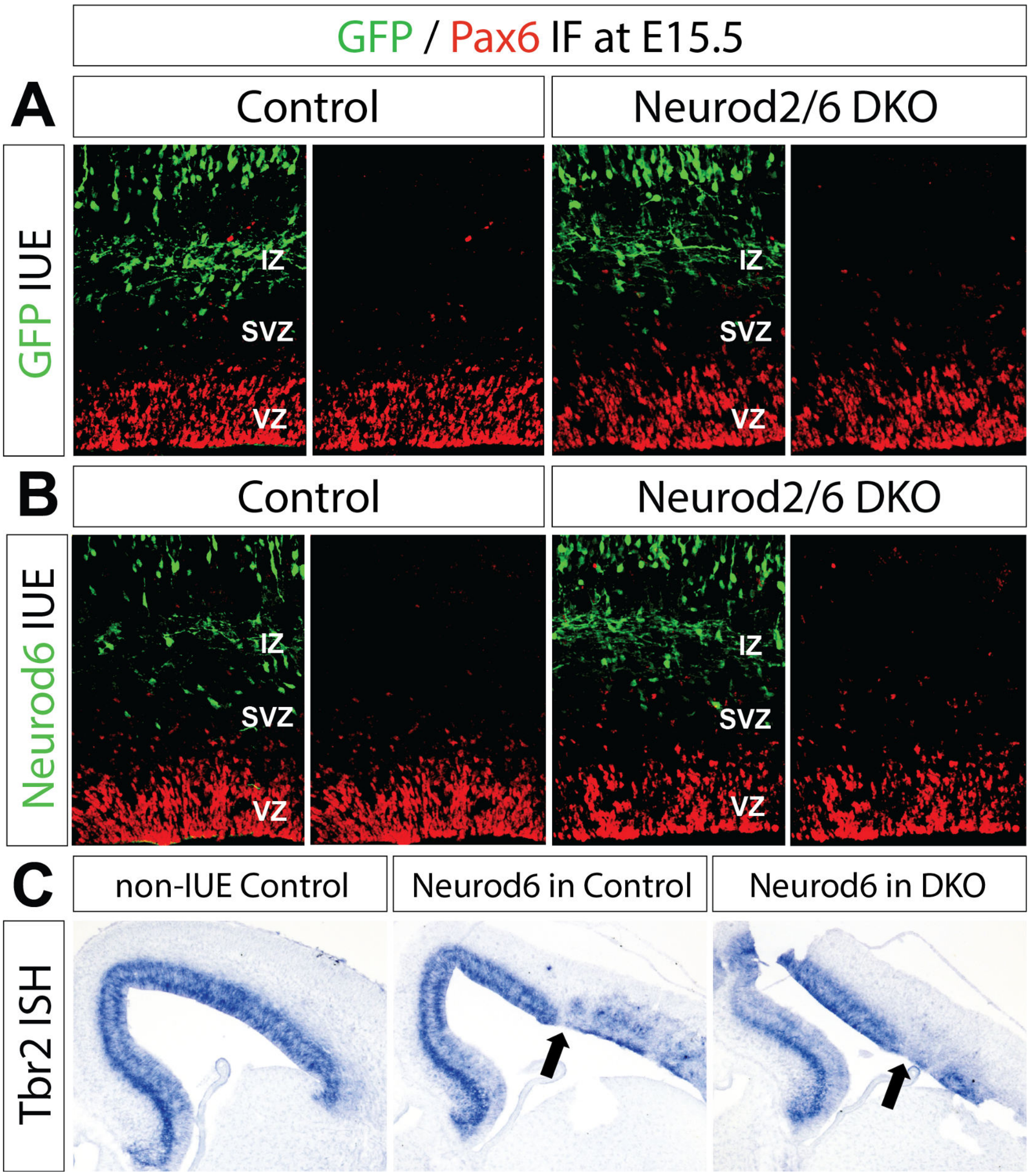


**Figure S6 Neurod6 over-expression does not disturb Pax6+ apical progenitors.**

**(A, B)** IF for GFP and Pax6 on GFP- or Neurod6-electroporated brain sections (at E15.5) shows that Neurod6 over-expression in both control and DKO brains does not disturb the development of Pax6+ APs.

**(C)** ISH for Tbr2 on Neurod6-electroporated brain sections. The brains were electroporated at E12.5 and fixed at E15.5. Tbr2 expression is downregulated at IUE sites.

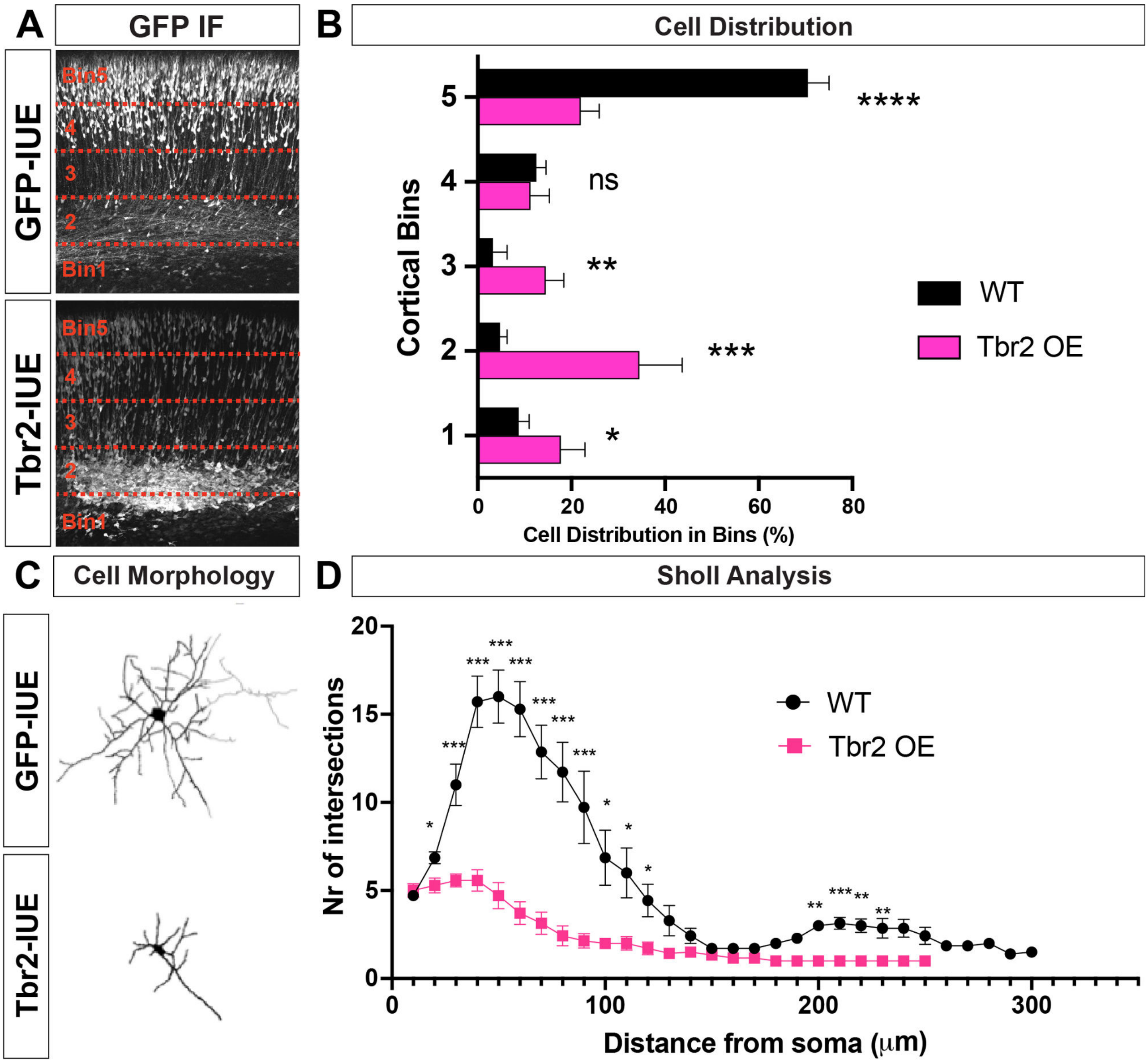


**Figure S7 Tbr2 over-expression impairs cortical neuron migration and neurite arborization**

**(A, B)** GFP and bicistronic Tbr2-GFP were electroporated into wild type (wt) brains at E14.5, which were then sampled at E18.5. IF for GFP shows that Tbr2 over-expression (OE) causes obvious retardation of neuron radial migration (**A**). The coronal brain sections were divided into 5 bins evenly and the cell proportions in each bin were quantified (**B**). GFP+ wt neurons follow normal radial migration, however, the cells carrying Tbr2 OE are largely restricted in deeper part of cortex: bin1 ($p$ = 0.032, *; n = 4), bin2 ($p$ = 0.00068, ***; n = 4), bin3 ($p$ = 0.0037, **; n = 4), but much less located in upper part: bin5 ($p$ < 0.0001, ****; n = 4).

**(C, D)** Cell morphology of GFP+ and Tbr2-electroporated cells in the cortex (**C**). (**D**) Stereotypical Sholl analysis shows neurite branching is severely curtailed in neurons carrying Tbr2 OE, particularly in the filed close to somas.

### Tables:

**Table S1 Primers**

| Name: | Usage: | Sequence: 5' – 3' |
|---|---|---|
| NeuroD2-Fw-EcoRI | Neurod2 full length cloning | TTTTGAATTCCAC-CATGCTGACCCGCCTGTTCAGC |
| NeuroD2-Rv-NotI | | TTTTGCGGCCGCTTATCAG-TTATGGAAAAATGCGTTG |
| NeuroD6-Fw-EcoRI | Neurod6 full length cloning | TTTTGAATTCAACCATGTTAACACTAC-CGTTTGAC |
| NeuroD6-Rv-NotI | | TTTT-GCGGCCGCTCATTAATTATGAAAAACTGCATTT |
| Tbr2-Fw-EcoRI | Tbr2 full length cloning | TTTTGAATTCTAAAGCATGCAGTTGG-GAGAGCAGC |
| Tbr2-Rv-NotI | | TTTTGCGGCCGCTCTAGGGACTTGTG-TAAAAAGCAT |
| Tbr2-ISH-Fw | Subcloning of Tbr2 cDNA fragment for ISH | AACAAGCTAGACATCGGTTCTTATG |
| Tbr2-ISH-Rv | | AATGCTCTAGGGACTTGTGTAAAAA |
| COUP-ISH-Fw | Subcloning of COUP-TF cDNA fragment for ISH | CCATAGATATGGCAATGGTAGTCAG |
| COUP-ISH-Rv | | AAGATCCGTATGTGGTCCATAAAA |
| Etv5-ISH-Fw | Subcloning of Etv5 cDNA fragment for ISH | GCTTCGCTTACTAAGTTTCTGAATG |
| Etv5-ISH-Rv | | GGAAGGGACTTGATTTACTTATGGT |
| Pax6-ISH-Fw | Subcloning of Pax6 cDNA fragment for ISH | ATGGAGAAGAGAAGAGAAACTGAG |
| Pax6-ISH-Rv | | ATACAAACTTGGAACATCAGTCCAT |
| Emx1-ISH-Fw | Subcloning of Emx1 cDNA fragment for ISH | AGAATCGGAGGACAAAATACAAACG |
| Emx1-ISH-Rv | | AGAACAAAGACAGAGACATGGAGA |
| Scrt2-ISH-Fw | Subcloning of Scrt2 cDNA fragment for ISH | GATCTTTCTTTTCATTTCTCAGCAG |
| Scrt2-ISH-Rv | | AAATACACAAATAAATAGCGGAAGG |
| Svet-ISH-Fw | Subcloning of Svet cDNA fragment for ISH | AAAGTCCTAATTAATGAAGATTGAG |
| Svet-ISH-Rv | | AGCTCCTTTACACTGAGCCTCTTCC |
| Brn1-ISH-Fw | | ATATATAACAAACAAAACCGGAAGA |

| | | |
|---|---|---|
| Brn1-ISH-Rv | Subcloning of Brn1 cDNA fragment for ISH | TATCGGAGTCAGTCCAGAAATAAAA |

**Table S2 Antibodies**

| Primary Anti-body | Species | Vendor | Cat # | Dilution | Usage | RRID |
|---|---|---|---|---|---|---|
| Neurod2 | Rabbit | Abcam | Ab104430 | IF: 1:500; Cut&Run: 1:50 | IF, Cut&Run | AB_10975628 |
| digoxigenin (AP-conju-gated) | sheep | Roche | 11093274910 | 1:1500 | ISH | AB_514497 |
| Tbr2 | Rabbit | Abcam | Ab183991 | 1:300 | IF | AB_2721040 |
| Tbr2 | Rat | Thermo Fisher | 14-4875-82 | 1:300 | IF | AB_11042577 |
| GFP | Goat | Rockland | 600-101-215 | 1:500 | IF | AB_828167 |
| GFP | Chicken | Abcam | Ab13970 | 1:1000 | IF | AB_300798 |
| BrdU | Rat | Abcam | Ab6326 | 1:500 | IF | AB_305426 |
| L1 | Rat | Millipore | MAB5272 | 1:1000 | IF | AB_2133200 |
| Pax6 | Rabbit | Millipore | Ab2237 | 1:500 | IF | AB_1587367 |
| Pax6 | Mouse | Thermo Fisher | MA1-109 | 1:500 | IF | AB_2536820 |
| Satb2 | Rabbit | homemade | - | 1:500 | IF | Britanova et al., 2008 |
| Ctip2 | Rat | Abcam | Ab18465 | 1:500 | IF | AB_2064130 |
| phospho-p44/42 ERK1/2 Thr202/204 | Rabbit | Cell Signal-ing | 9101 | 1:200 | IF | AB_331646 |
| Brn2 | Goat | Santa Cruz | sc-6029 | 1:200 | IF | AB_2167385 |
| Tbr1 | Rabbit | Abcam | ab31940 | 1:300 | IF | AB_2200219 |

| | | | | | | |
|---|---|---|---|---|---|---|
| Sox5 | Goat | Santa Cruz | sc-17329 | 1:200 | IF | discontinued |
| Oct6 | Rabbit | Abcam | ab272925 | 1:300 | IF | AB_2927579 |
| Foxp2 | Rabbit | Abcam | ab16046 | 1:500 | IF | AB_2107107 |
| Cre | Mouse | Chemicon | MAB3120 | 1:200 | IF | AB_2085748 |
| Nestin | Mouse | Millipore | MAB353 | 1:500 | IF | AB_94911 |
| BLBP | Rabbit | Millipore | ABN14 | 1:1000 | IF | AB_11210770 |
| Reelin | Mouse | R&D | AF3820 | 1:500 | IF | AB_2253745 |
| PCNA | Mouse | Santa Cruz | sc-56 | 1:200 | IF | AB_628110 |
| Ki67 | Rabbit | Abcam | ab15580 | 1;500 | IF | AB_443209 |
| Sox2 | Mouse | Santa Cruz | sc-365823 | 1:200 | IF | AB_10842165 |
| Olig2 | Rabbit | gift of Prof. C. Stiles | - | 1:500 | IF | Heather et al., 2004 |
| Tuj1 | Mouse | Covance | MRB-435P | 1:1000 | IF | AB_663339 |
